# m^6^A positions polyadenylation in *Plasmodium falciparum*

**DOI:** 10.64898/2026.05.19.726191

**Authors:** Joshua M. Levendis, Amy Distiller, Selina Mussgnug, Lakvin Fernando, Steven Lancashire, Neil D. Young, Emma McHugh, Sebastian Baumgarten, Stuart A. Ralph

**Affiliations:** Department of Biochemistry & Pharmacology, Bio21 Molecular Science and Biotechnology Institute, The University of Melbourne; Institut Pasteur, Université Paris Cité, INSERM U1347, G5 Parasite RNA Biology Group, F-75015 Paris, France; Department of Veterinary Biosciences, Melbourne Veterinary School, Faculty of Science, The University of Melbourne, Parkville, VIC 3010, Australia

## Abstract

N^6^-methyladenosine (m^6^A) is the most abundant modification in eukaryotic mRNA and can influence gene expression, but the molecular mechanisms by which it does this at an individual gene level are poorly understood. The eukaryote with the highest known percentage of adenosine in its RNA is *P. falciparum*, causative agent of the deadliest form of malaria. *P. falciparum* has a high genomic AT content (>80% A+T) and a particular A-bias in mRNA - coding strands have an average adenosine content of 44%. *Plasmodium* parasites employ extensive transcriptional and post-transcriptional control over gene expression to differentiate within and between sexual and asexual stages in response to changing environments and hosts. We used direct Nanopore RNA sequencing and GLORI-seq to interrogate m^6^A methylation of the asexual *Plasmodium falciparum* transcriptome, revealing m^6^A depletion in protein coding regions relative to the 3’UTR, where m^6^A is deposited in precise patterns approximately 50 nt upstream of the polyadenylation site. We used an inducible protein mislocalisation system to disrupt the methyltransferase, which writes m^6^A to mRNA, and observed m^6^A depletion associated with transcriptional readthrough and chimeric transcripts. We show that m^6^A in transcript 3’UTRs is required for faithful positioning of polyadenylation and transcription termination. This work highlights the importance of m^6^A to mRNA processing in *P. falciparum*, and potentially a wider role in post-transcriptional regulation.

## Introduction

*Plasmodium* is the causative agent of malaria – a bloodborne infectious disease. Malaria is present in 40% of the world and kills 400,000 people every year, 80% of whom are children (1). There are over 200 known species of *Plasmodium* that infect reptiles, birds and mammals, yet only six regularly infect humans (*Plasmodium falciparum*, *P. ovalecurtisi, P. ovalewallikeri*, *P. malariae*, *P. vivax*, *P. knowlesi*) with *P. falciparum* causing the most severe symptoms and deaths in humans (2). *Plasmodium* parasites need a strict control over gene expression to permit their multiple distinct stages and morphology. The *Plasmodium falciparum* genome is small (23 Mbp, ∼5400 protein coding genes) and has a high A+T content compared to other organisms (∼80% AT pairs in coding and 90% in intergenic regions) (3). In addition to a nuclear genome, *Plasmodium* has transcription from two organellar genomes, both also AT rich: an apicoplast genome (35 kbp circular genome, 30 protein coding genes), and a mitochondrial genome (6 kb linear genome with 3 protein coding genes). In both organelles transcription is polycistronic, and the apicoplast contains its own splicing and RNA modification machinery (4, 5). Nuclear transcription is monocistronic like most other eukaryotes. *Plasmodium* employs both transcriptional and post-transcriptional control over gene expression. For example, anti-sense regulation of some genes appears to initiate gametocytogenesis (6, 7), while post-transcriptional silencing of other transcripts is key to gametogenesis (8).

One source of post-transcriptional control in eukaryotes is RNA modification, which can influence mRNA fates by changing RNA base-paring (*cis*) or via effector proteins that recognise specific modifications (*trans*) (9, 10). This in turn influences diverse RNA fates, including splicing, mRNA processing, stability, trafficking and translational efficiency (11). Multiple RNA modifications have been identified in *Plasmodium,* but the exact molecular mechanisms by which they exert influence are still poorly understood (9, 12–14). The primary pathway by which RNA nucleosides are modified involves families of writer and eraser enzymes which bind to RNA co- or post-transcriptionally and add or remove functional groups (15–20). These enzymes can have site-specific activity, leading to ‘canonical modifications’ which consistently appear in RNA locations determined by sequence or structure (21). Writer enzymes can also have non-specific activity, resulting in sporadic or seemingly random modifications which are not conserved across RNA types (16). Modified RNA nucleotides can also be randomly incorporated by RNA polymerases, which are more error prone than DNA polymerases (22–24).

We are interested in how the most abundant eukaryotic mRNA modification, N^6^-methyladenosine (m^6^A), affects *P. falciparum* gene expression. Due to its high AT content, we speculate that there are elevated levels of m^6^A compared with other organisms. m^6^A is written to nascent mRNA transcripts by the m^6^A writer complex. In mammals, this complex consists of the catalytic component (METTL3, METTL14) and a collection of proteins which regulate and facilitate RNA binding (WTAP, HAKAI, VIRMA, ZC3H13, and RBM15) (11). *P. falciparum* has orthologues of METTL3, METTL14 and WTAP (named *Pf*MT-A70, *Pf*MT-A70.2 and *Pf*WTAP respectively), yet co-immunoprecipitation of the *P. falciparum* methyltransferase complex showed a total of at least 14 additional interacting proteins, including nucleotide binding proteins (9). The m^6^A writer complex in eukaryotes preferentially binds and writes to adenosines within the DRACH motif (D=A/G/U, R=A/G, H=A/C/U) (25, 26), yet the presence of this motif does not guarantee methylation (27). The DRACH motif occurs throughout coding and untranslated regions of mRNA but m^6^A occurs preferentially around stop codons and 3’UTRs (28). In *Toxoplasma* and *Arabidopsis,* m^6^A acts as a guard to prevent transcriptional readthrough, and in mammals, perturbation of the m^6^A eraser changes mRNA length (29, 30). While a previous m^6^A pull down identified genes preferentially modified with m^6^A in *P. falciparum* (9), a base-level high resolution methylome does not exist and it is unknown to what extent m^6^A controls mRNA length.

We can study m^6^A in *P. falciparum* by perturbing the m^6^A writer and reader enzymes (9, 12, 31). In this study we employ a knock-sideways system to inducibly mislocalise the *P. falciparum* writer subunit *Pf*MTA-70 (PF3D7_0729500) from the nucleus to the parasite plasma membrane. We use Oxford Nanopore Technologies (ONT) direct RNA sequencing (SQK-RNA004) to quantify gene expression and detect RNA modifications at different timepoints just prior to *P. falciparum* asexual replication (schizogony) in parasites with *Pf*MTA-70 mislocalised and untreated parasites. *Pf*MTA-70 perturbation knocked down m^6^A modifications in mRNA and led to changes in both overall gene expression and in transcript length. Specifically, we found that m^6^A modifications are enriched in transcripts approximately 50 nt upstream of the site of 3’ cleavage and polyadenylation. Depleting m^6^A at these positions increased average 3’UTR length and caused an increase in chimeric mRNA abundance. Our study reveals a role for m^6^A in defining the site of 3’ processing of mRNA, which is faulty in the absence of that modification.

## Results

### A knock-sideways system mislocalises the m^6^A methyltransferase

We studied how m^6^A regulates mRNA in asexual parasites by harvesting late-trophozoites (Figure 1A). In *P. falciparum,* m^6^A is written to nascent mRNA by the methyltransferase complex (Figure 1B). The proteins in this complex which share homology with other eukaryotes are *Pf*MT-A70, *Pf*MT-A70.2 and *Pf*WTAP. We tagged the catalytic subunit *Pf*MT-A70 with a 2xFKBP-GFP-2xFKBP C-terminus (hereafter referred to as *Pf*MT-A70-sandwich) and transfected an exogenous pLyn-FRB-mCherry plasmid to act as a parasite plasma membrane mislocaliser (32). We confirmed that our transfectants had the sandwich construct integrated correctly into the relevant locus, and created the desired fusion transcript, through RNA-seq (Figure 1C). Upon addition of rapamycin, the FKBP and FRB domains dimerise, forcing mislocalisation of the methyltransferase to the parasite membrane where it cannot effectively methylate nascent transcripts (Figure 1D). The inducible sequestration of *Pf*MTA-70 allows temporal study of m^6^A in *P. falciparum,* important for studying stage specific gene control. We observed mislocalisation of the *Pf*MT-A70-sandwich GFP signal at 28, 32 and 36 hours post invasion (hpi) after 4 hours on rapamycin, as indicated by the shift of condensed to diffuse green fluorescence (Figure 1E). *Pf*MT-A70 mislocalised parasites demonstrate a noticeable growth-defect over a single cycle compared to parasites that were not treated with rapamycin (Figure 1F) or compared to parental 3D7 strain parasites that were treated with rapamycin. However, this *Pf*MT-A70 mislocalisation did not lead to growth collapse – parasites were able to grow continuously (though slightly more slowly) for multiple cycles even with continuous rapamycin mislocalisation of *Pf*MT-A70.

**Fig 1.**
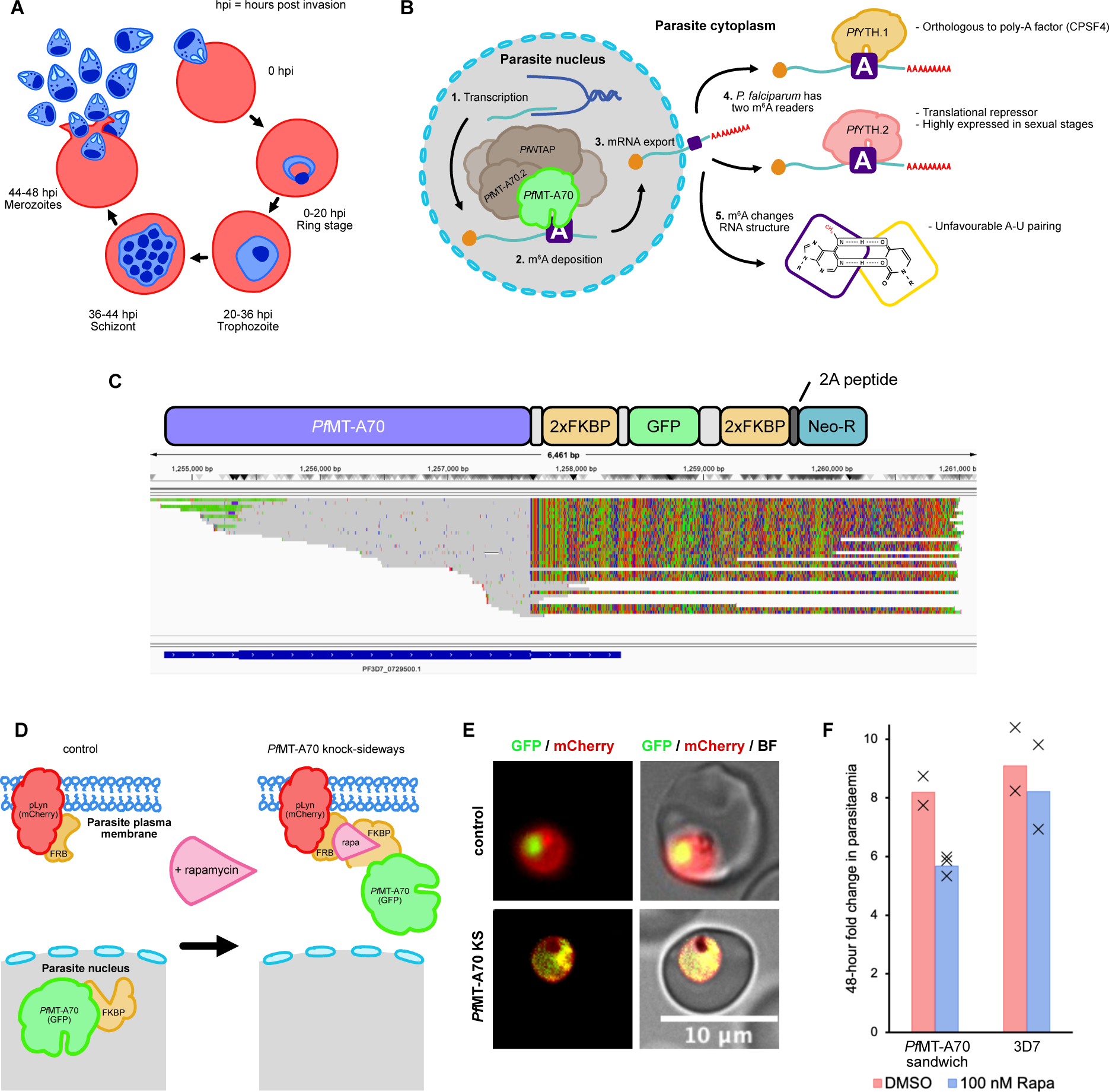
A knock-sideways system mislocalises the m^6^A methyltransferase. **A)** *Plasmodium falciparum* has 48 hour lifecycle in asexual blood stages. **B)** m^6^A is deposited to nascent mRNA in the nucleus by the methyltransferase complex, of which three subunits share homology with humans (*Pf*WTAP, *Pf*MT-A70, *Pf*MT-A70.2). m^6^A can influence mRNA fate in the cytoplasm by interacting with m^6^A readers (YTH domain containing proteins) or altering mRNA secondary structure. **C)** *Pf*MT-A70-sandwich construct design (top) aligned to IGV screenshot of ONT RNA-seq (bottom). Bases that aligned correctly to *Pf*MT-A70 locus are shown in grey and mismatched in colour (A=green, T=red, C=blue, G=orange). Mismatched bases represent knock-sideways domains inserted at the 3’ end of CDS. **D)** A knock-sideways system can be used to mislocalise the methyltransferase subunit *Pf*MT-A70 from the nucleus to the parasite plasma membrane. Upon addition of the mTOR inhibitor rapamycin, a FKBP-FRB heterodimer is formed around rapamycin. **E)** Mislocalisation can be verified using fluorescence microscopy. Late-trophozoites are shown (32 hpi) with 4 hours on rapamycin (bottom). **F)** Growth assays for *Pf*MT-A70-sandwich and *Pf*3D7 challenged with 100 nM rapamycin.

### Nanopore direct RNA sequencing allows study of modifications and expression levels at the same time

To study how *Pf*MT-A70-sandwich mislocalisation changed m^6^A we performed direct RNA nanopore sequencing at different timepoints in the late-trophozoite stage and filtered for high quality transcripts (Supplementary Table 1). Our ONT RNA-seq was able to quantitatively detect the RNA modifications m^6^A, inosine (I), pseudouridine (Ψ) and N^5^-methylcytidine (m^5^C) using the nanopore modified basecaller Dorado (33) (Figure 2A, Supplementary Figure 1). All the above 4 modifications were detected in *P. falciparum* mRNA, with m^6^A the most abundant modification transcriptome-wide. Different RNA modifications had different positional biases in transcripts; m^6^A was significantly enriched at specific sites in the 3’UTR, Ψ and I appeared scattered throughout the transcript body with no discernible pattern from visual inspection, and m^5^C was lowest in abundance of the modifications detected and appeared enriched in coding sequences. The latter distribution is plausibly connected to the relative GC richness in *P. falciparum* coding regions relative to UTRs (20% vs 10% GC content).

**Fig 2.**
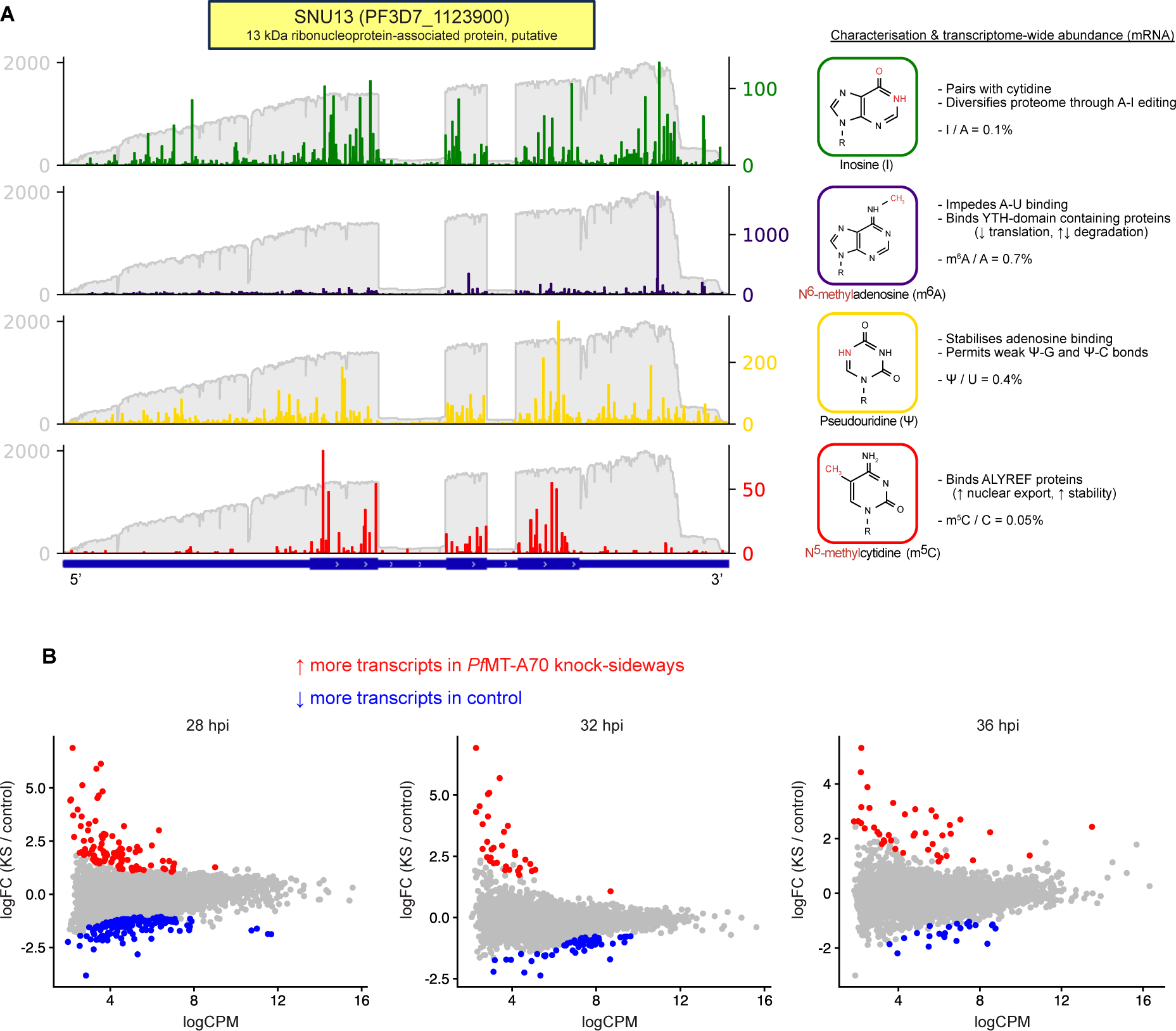
Nanopore direct RNA sequencing enables studying modifications and expression levels at the same time. **A)** Coverage plots show modification coverage of example gene (SNU13) using ONT SQK-RNA004, accompanied by descriptions and prevalence statistics for modifications that Nanopore can detect. Note the changing left and right y-axes which represent the differential abundance of each modification. **B)** edgeR differential expression (DE) analysis shows comparable levels of genes with increased and decreased transcript counts after *Pf*MT-A70 knock-sideways (KS), with no GO terms enriched (Benjamini-Hochberg FDR < 0.05, GLM likelihood ratio test).

We speculated that since m^6^A is widely reported to be involved in RNA destabilisation, that m^6^A depletion would increase transcript stability. However, we found *Pf*MT-A70-sandwich mislocalisation caused an even split of increased and decreased abundance of all transcripts (Figure 2B, Supplementary Tables 2-5). Principal component analysis of gene expression counts showed samples separated by timepoint but not treatment (Supplementary Figure 2). This indicates that our three timepoints harvested had distinct expression levels (consistent with known temporal regulation of transcript regulation in *P. falciparum*) but suggests expression changes induced by m^6^A depletion were not a consistent separating factor.

### Nanopore can detect m^6^A depletion after *Pf*MT-A70 knock-sideways

Although *Pf*MT-A70 knock-sideways appeared to have little short-term impact on mRNA abundance, mislocalisation of this methyltransferase had a rapid impact on the degree of m^6^A methylation of transcripts (Figure 3A). When we manually inspected genes and their aligned transcripts in Integrative Genomics Viewer (IGV) we observed clear differences in m^6^A methylation. This was true both for m^6^A that occurred at sites with frequent and consistent methylation, as well as for positions with sporadic methylation (Figure 3A, Supplementary Figure 3A). Hereafter, we will refer to transcript base positions with a high ratio of m^6^A methylation (m^6^A / A ≥ 50%) as ‘canonical’ sites and others as ‘low m^6^A-frequency.’

**Fig 3.**
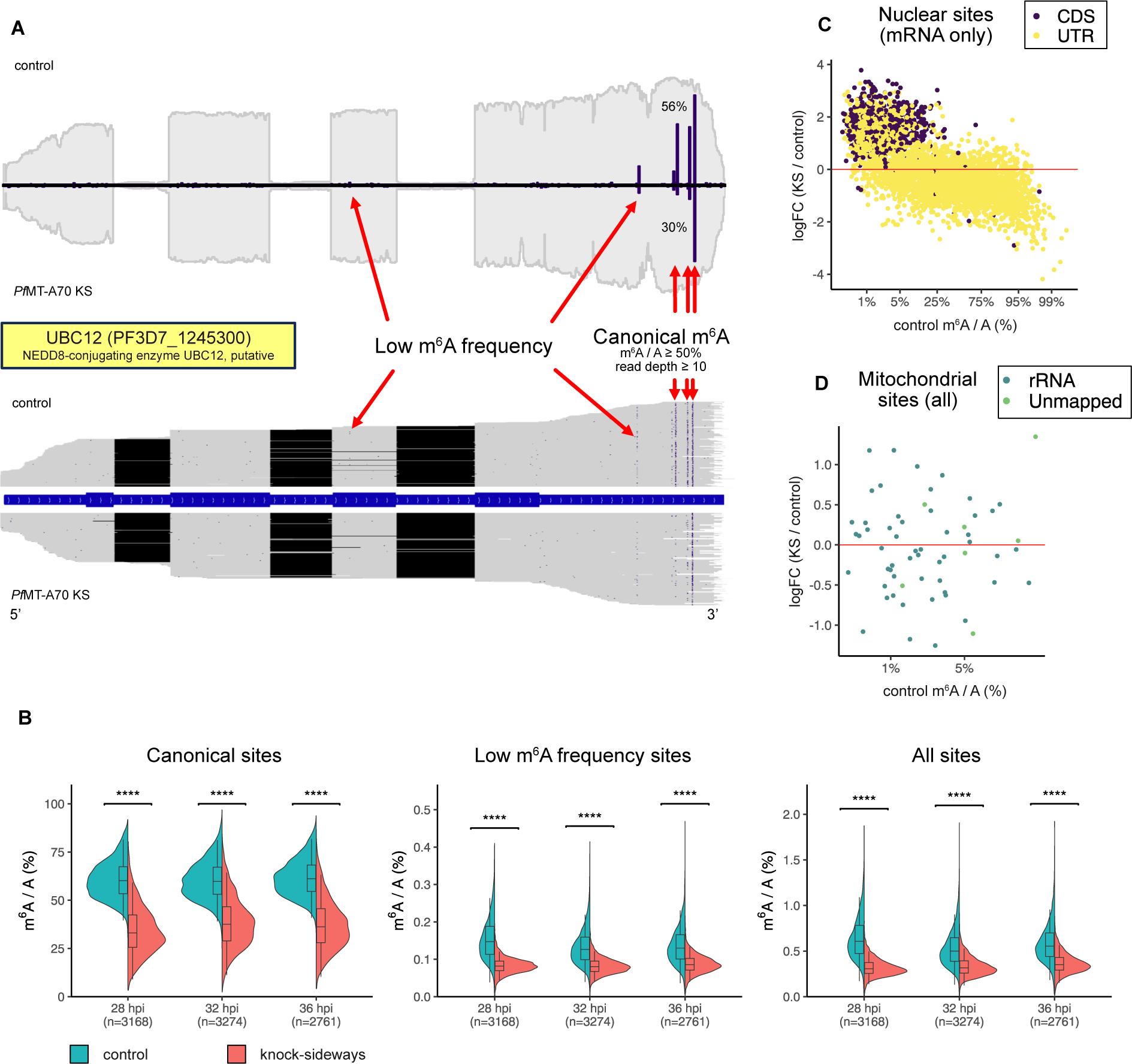
Nanopore can detect m^6^A depletion after *Pf*MT-A70 mislocalisation. **A)** Example gene coverage (UBC12) showing transcript level m^6^A depletion. IGV screenshot (bottom) shows sites within individual transcripts. Sites are considered canonically methylated (m^6^A / A ≥ 50%) or low frequency (m^6^A / A ≤ 50%). Weighted average methylation (WAM) of canonical m^6^A sites is shown alongside gene coverage, where each m^6^A site is weighted against its read depth. This metric can be applied to canonical m^6^A, low frequency, or all sites within each gene to provide gene level methylation analysis. **B)** Quantification of gene-level WAM for canonical sites, low frequency sites, and total m^6^A / A before and after *Pf*MT-A70 knock-sideways in *P. falciparum* mRNA (28, 32 and 36 hpi). After *Pf*MT-A70 mislocalisation, both canonical and low-frequency sites have reduced methylation (paired t-test, p < 0.0001). **C)** Site level differential methylation analysis shows nuclear encoded individual m^6^A sites are differentially methylated after *Pf*MT-A70 knock-sideways but **D)** mitochondrial sites are not (28 hpi, 4-hour knock-sideways, edgeR GLM likelihood ratio test). Sites with low methylation are more likely to increase in methylation after knock-sideways.

We observed that genes and their transcripts often contained multiple canonical m^6^A sites. Most genes contained one or two canonical m^6^As, with a maximum detected of seven canonical sites (Supplementary Figure 4E). To quantify methylation changes at the gene level we calculated average canonical m^6^A changes weighted with respect to the read depth of each gene. We quantified methylation rate with and without knock-sideways for these sites (averaged at the gene level) and found while low m^6^A frequency sites were far less methylated on average than canonical sites (0.1% vs 60%) they both had a significant reduction in m^6^A after knock-sideways (Figure 3B, Supplementary Tables 6-8). With these gene level methylation rates, we performed a PCA and found that samples separated strongly by treatment (80% PC1), and weakly by timepoint (4% PC2) (Supplementary Figure 4A).

We then queried what differential methylation looked like at the site level and found the individual sites with very low initial methylation rates tended to have an increase in m^6^A upon knock-sideways (Figure 3C). This was counterintuitive to what we expected but could be explained as statistical artifacts from increased mRNA length. We found that sites in coding regions tended to have lower methylation (< 25%) and highly methylated sites were almost entirely in UTR regions. Even though our ONT library preparation selects for polyadenylated RNAs, we detected non-coding RNAs including rRNA, snoRNA and few tRNAs (Supplementary Figure 3, 5). Non-coding RNA sites appeared to increase in methylation rate after knock-sideways in a similar fashion to the coding region (Supplementary Figure 3B). Often transcripts would extend beyond the annotated gene boundaries but were still methylated, and we found these ‘unmapped’ sites followed a similar methylation pattern to the mRNAs. We found that mitochondrial sites were lowly methylated (< 10%) and not significantly differentially methylated after knock-sideways (Figure 3D). This is consistent with anticipated separate mechanisms for modification of organellar and nuclear-encoded RNAs. We also observed positional enrichment of pseudouridine, m^5^C and m^6^A in tRNAs and rRNAs which conincided with known modification positions in these RNAs in other eukaryotes (34, 35), as well as modifications in mitochondrial genome-encoded genes (Supplementary Figure 6). Although the accurate detection of known modifications at these positions on other RNA molecules is a reassuring validation of our detection, the scope of this paper is limited to m^6^A in nuclear-encoded mRNA, and we limit our analysis below to these transcripts.

### m^6^A is enriched at a conserved DRACH-like motif

In parallel to analysing the m^6^A methylome by ONT, we performed GLORI-seq on *P. falciparum* RNA at 12, 24 and 48hpi from asexual blood stage parasites (36). GLORI-seq is an alternative sequencing method that detects m^6^A by quantifying m^6^A as errors in reverse transcription (Figure 4A). Since there is no ground truth methylome for *P. falciparum*, we compared data of our canonical sites between ONT and GLORI-seq data (Supplementary Tables 9-10). These datasets were produced at different read depths and are likely to have different signal:noise ratios. We therefore opted to use a lower threshold for this comparison only for canonical m^6^A detection as determined by the GLORI-seq dataset (m^6^A / A ≥ 25%, read depth ≥ 15).

**Fig 4.**
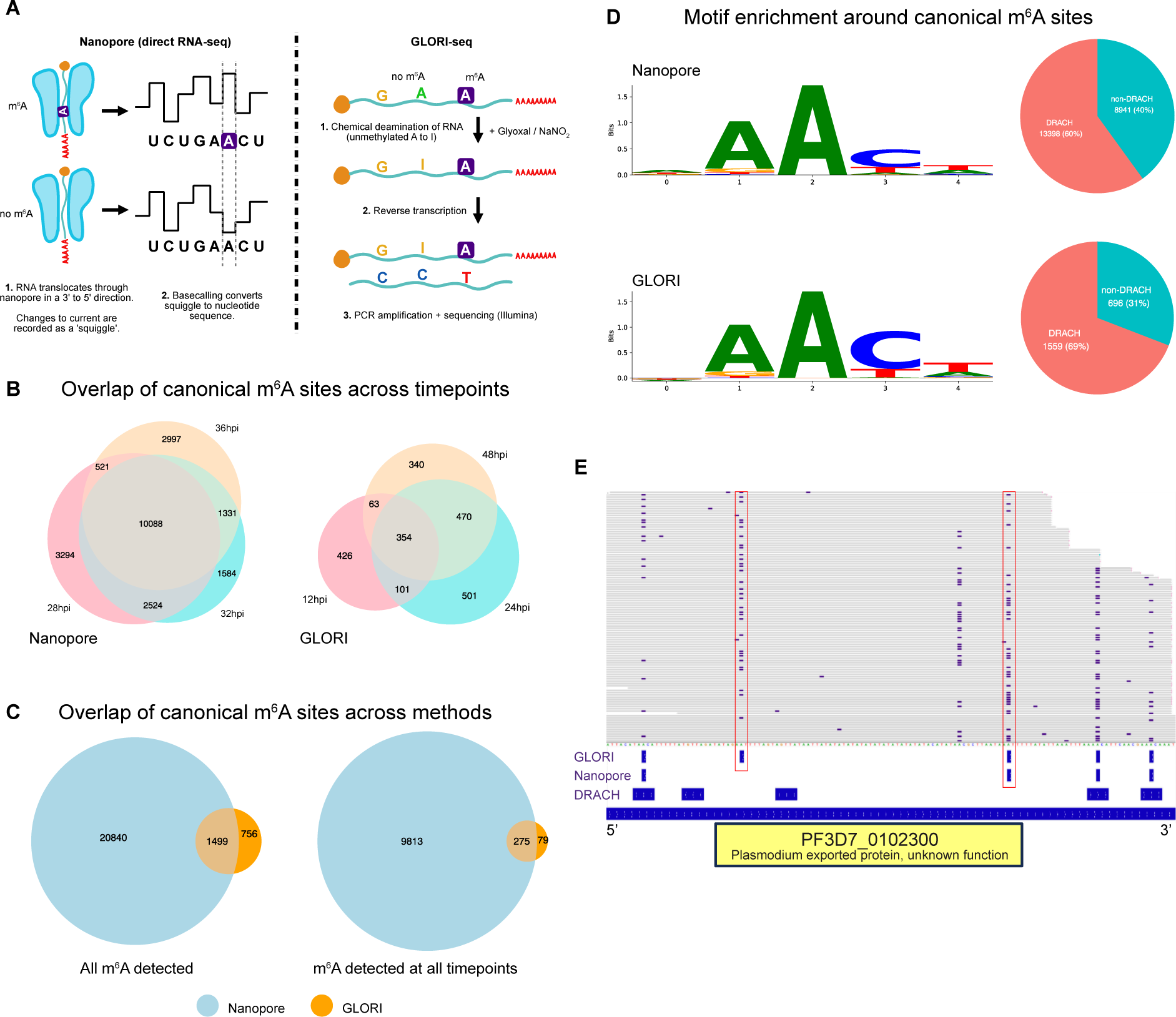
m^6^A sites detected by Nanopore and GLORI-seq are enriched at a conserved motif. **A)** Schematic of Nanopore vs GLORI modification sequencing methods. **B)** Canonical sites detected by Nanopore and GLORI at different timepoints. Here we define canonical sites as m^6^A / A ≥ 25%, depth ≥ 15. **C)** Canonical sites detected by both methods for all m^6^A detected (left) or sites only detected at all timepoints (right). **D)** Sequence logo showing conserved genomic motifs where m^6^A is detected in *P. falciparum* mRNA. Canonical m^6^As occur both at the degenerate consensus motif DRACH (∼60%) and non-DRACH sites, in both Nanopore and GLORI experiments (D=A/G/U, R=A/G, H=A/C/U). **E)** IGV screenshot (cropped) showing Nanopore and GLORI agreement on methylated sites, and non-DRACH sites (in red boxes).

To account for differences in timepoints sequenced, we decided to compare only high confidence m^6^A sites which were present at all three timepoints in each experiment, 10,088 in ONT vs 354 in GLORI-seq (Figure 4B). 78% of canonical m^6^A were detected by both methods, with ONT detecting far more sites overall (Figure 4C). Different read depth between experiments (GLORI-seq ∼300 read depth, ONT ∼100,000), plausibly explains the discrepancy in total sites detected.

Both experiments identified similar conserved DRACH sequence motifs that were favourably methylated and detected similar proportions of non-DRACH methylation (Figure 4D, E).

### m^6^A position predicts the position of polyadenylation sites

We compared m^6^A gene body position between GLORI-seq and nanopore experiments. Both methods detected preferential 3’UTR distribution of canonical m^6^A, and both methods observed a bimodal peak around the end of the 3’UTR (Figure 5A). The bimodal peak is likely an artefact of imprecise gene annotation, as we often observed gene annotations ended before or after the true 3’ end of transcripts many genes. We observed a slight increase in m^6^A abundance in and upstream of the annotated 5’UTR. Again, we think this is more likely an artefact of poor gene annotations and represents m^6^A in the 3’ of transcripts for adjacent upstream neighbour genes where mapping assigned a read to the inappropriate gene.

**Fig 5.**
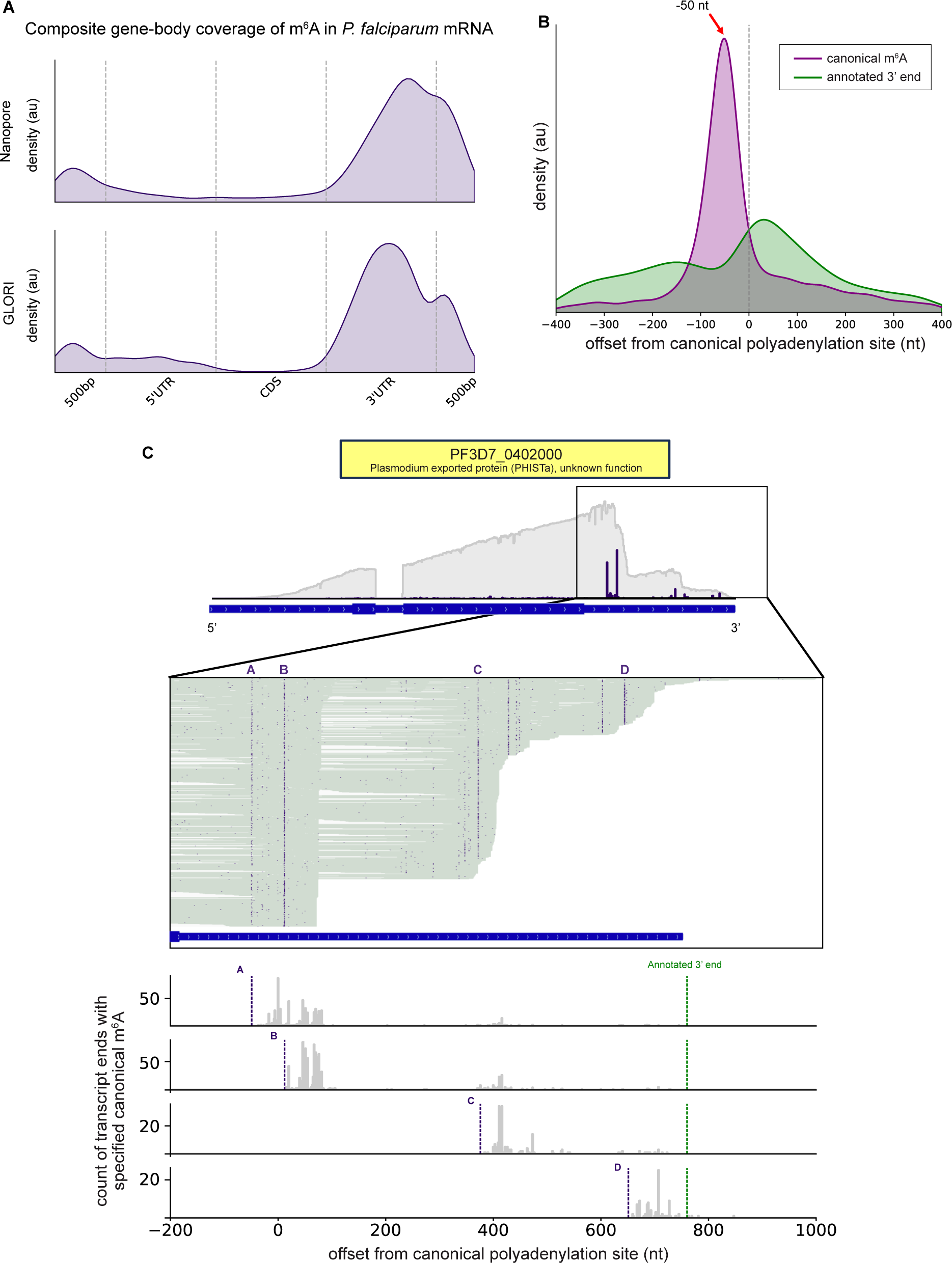
m^6^A is enriched near polyadenylation sites. **A)** Composite gene-body coverage shows canonical m^6^A are primarily found in 3’UTRs in *P. falciparum* mRNA, similar to other eukaryotes. Here, we define canonical m^6^A as sites with m^6^A / A ≥ 50%. **B)** Canonical m^6^As occur close (∼50 nt) to polyadenylation sites. We define canonical polyadenylation site as the most common polyadenylation site for all transcripts aligned to a gene. **C)** Example gene containing multiple canonical m^6^As (lettered), and consequently multiple polyadenylation sites (Cropped IGV screenshot). Genomically aligned RNA transcripts are shown as horizontal lines (green) and contained m^6^A as purple squares. Bottom panel shows histogram of individual transcript polyadenylation sites, separated by which m^6^A they contain.

We endeavoured to measure the average distance of m^6^A from the polyadenylation site in 3’UTRs in our ONT RNA-seq, while accounting for annotation errors. We approximated ‘canonical’ polyadenylation sites by taking the genomic position for each gene with the most transcript end / poly-A tail start sites (Supplementary Tables 11-13). We then plotted for each gene the distance of canonical m^6^As and 3’ annotation end against this offset and found that, on average, m^6^As are ∼50 nt upstream of polyadenylation sites (Figure 5B). We observed a similar bimodal peak in the annotation end offsets as seen in the gene body distribution plots (with reversed 5’ **→** 3’ directionality), but not in the overall m^6^A distribution, confirming that the bimodal peak is due to gene annotation errors.

Often, genes have multiple canonical m^6^As (Figure 3A, Figure 5C, Supplementary Figure 4E, F). We asked how the m^6^A-poly-A distance of ∼50 nt changes for genes with complex 3’UTR methylation. With ONT RNA-seq we were able to separate transcripts by which canonical m^6^A they contain and quantify their polyadenylation sites. In this case, each site of canonical methylation has a polyadenylation site roughly ∼50 nt downstream (Figure 5C), showing that for genes with complex methylation, each canonical m^6^A is usually associated with one alternative polyadenylation site.

Finally, we asked if transcripts that have m^6^A are more likely to terminate within 100nt than unmethylated transcripts. For each canonical m^6^A in our data, we separated transcripts into subpopulations based on whether they were methylated or not at the given site, and calculated the probability that transcripts would terminate within 100 nt (Figure 6A). Indeed, we found that m^6^A was a strong predictor of polyadenylation within that 100 nt window in both the control and knock-sideways treatments (Figure 6B, 6C). Interestingly, when comparing the probability of proximal polyadenylation in only methylated transcripts between control and knock-sideways there was no significant difference, yet comparing unmethylated transcripts between treatments showed a change. This suggests that m^6^A methylation guides polyadenylation similarly regardless of treatment, but unmethylated transcripts are not guaranteed to terminate in a similar fashion across treatments.

**Fig 6.**
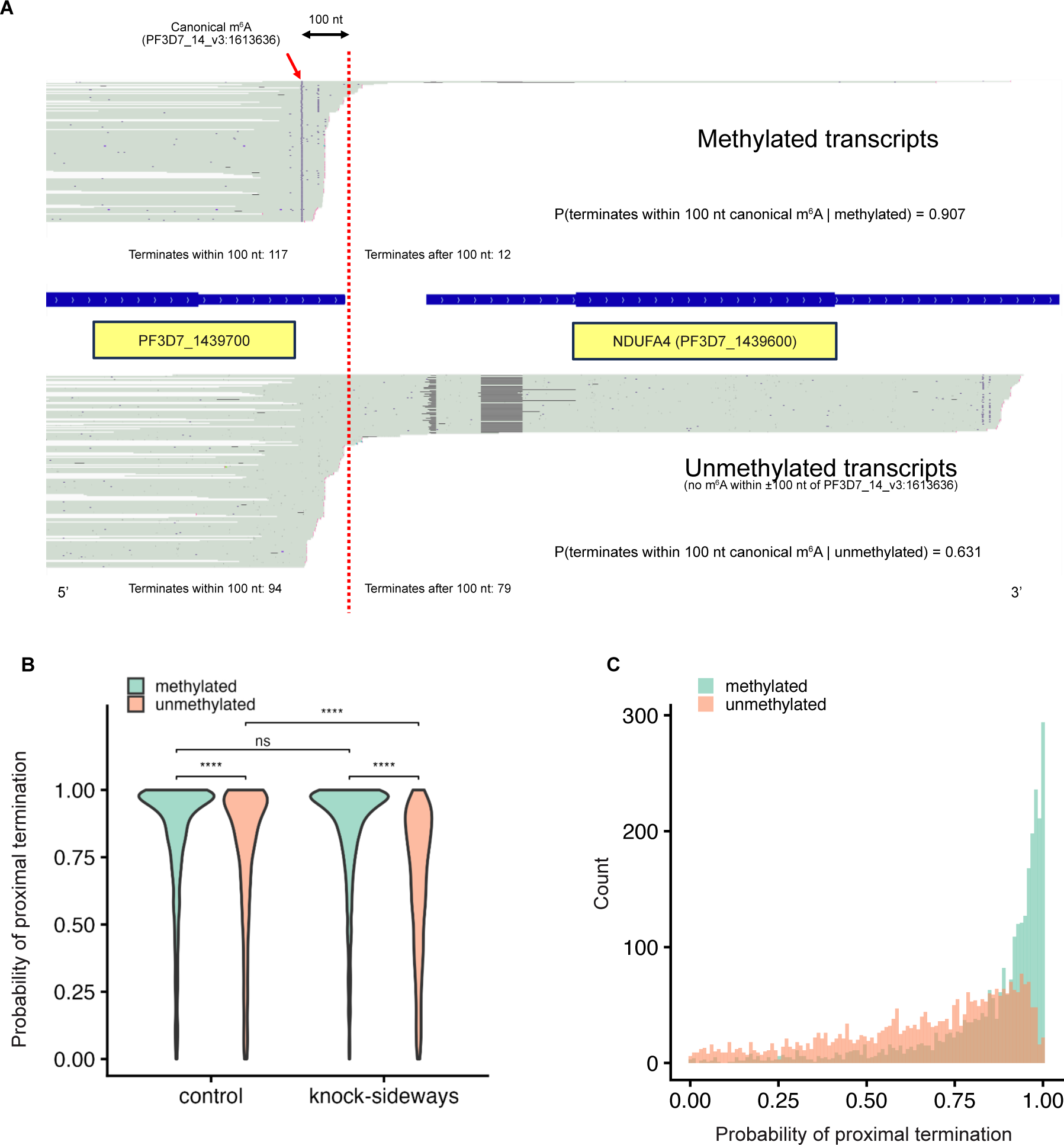
Single molecule analysis shows methylation status is a predictor of proximal transcript termination. **A)** Grouping individual transcripts by methylation status within a sample (28K1). IGV screenshots show example separation of transcript subpopulations based on methylation at a given site (annotated with red arrow) into methylated or unmethylated groups (no m^6^A within ±100 nt of a given m^6^A site). From these subpopulations, the probability that transcripts containing the methylated site will terminate within 100 nt can be calculated. In this example, we show methylated transcripts have a higher probability of proximal termination. **B)** Distribution of proximal termination probabilities for all m^6^A sites in transcriptome (2915 sites with ≥ 50% m^6^A / A). Methylated transcripts have a high probability of terminating within 100 nt of a canonical m^6^A regardless of condition (ns, two-sided wilcox test). Unmethylated transcripts are more likely to run on when missing an m^6^A. **C)** Histogram showing binned count of probabilities in knock-sideways treatment (bin width = 0.01).

### *Pf*MT-A70-sandwich knock-sideways disrupts normal transcript termination and increases chimeric transcript abundance

*P. falciparum* has two described m^6^A readers – *Pf*YTH.1 and *Pf*YTH.2 (9). *Pf*YTH.1is orthologous to CPSF4 / CPSF30 (37), which complexes with transcriptional cleavage and polyadenylation machinery (CPSF). In *Toxoplasma* and *Arabidopsis*, the knockout of this m^6^A reader, or methyltransferase machinery changes 3’UTR length (30). Indeed, disruption of methyltransferases as well as demethylases in mammalian cells affects transcript length (29). We therefore asked how UTR length changed when *Pf*MT-A70 was mislocalised. We saw a marginal increase in average UTR length upon *Pf*MT-A70 mislocalisation (Supplementary Figure 7A). The moderate UTR length change is likely due to only a small proportion of a genes total transcripts having increased UTR length. Using our canonical polyadenylation approximation described above, we also quantified the proportion of run-on transcripts – those which terminate beyond the usual polyadenylation site – and detected an increase in number of run-on transcripts after knock-sideways (Supplementary Figure 7B), but no consistent change in polyA length (Supplementary Figure 7C). The spread of run-on transcripts even before m^6^A depletion can be attributed to the difficulty in estimating a point to define run-on transcripts in genes with multiple canonical polyadenylation sites. We attempted to correlate UTR changes and run-on proportion changes with m^6^A depletion but found only weak correlation (Supplementary Figure 7D). We suspect that the existence of multiple canonical m^6^As in many transcripts confounds our ability to identify a clearer relationship between loss of m^6^A and increased run-on.

In inspecting the genes that were identified as most differentially expressed after *Pf*MT-A70 knock-sideways we recognised that several of these fell into a specific category. Transcripts for these genes were barely detected or not detected at all in the untreated parasites, but were flagged as being expressed after *Pf*MT-A70 mislocalisation. Visual inspection in IGV revealed that these newly activated genes were in fact the result of an adjacent, upstream gene whose transcription had aberrantly continued through into the downstream, otherwise unexpressed gene (Supplementary Figure 8E). We classified these transcripts that included multiple protein coding genes as chimeric transcripts, and we specifically investigated if such chimeric transcripts showed a global change after m^6^A depletion. Such chimeric transcripts likely result from failed transcriptional termination of the upstream gene after m^6^A depletion. We considered chimeric transcripts as those which mapped to more than one gene by featureCounts (38) (ambiguous transcripts). We acknowledge this is less stringent than filtering for only transcripts which have introns from multiple genes spliced, however this approach also highlights transcripts which map to downstream UTRs, which may still impact expression. We saw an overall increase in chimeric transcripts after m^6^A depletion as revealed by differential abundance analysis of chimeric transcripts with respect to all transcripts (Figure 7A-C, Supplementary Tables 14-17).

**Fig 7.**
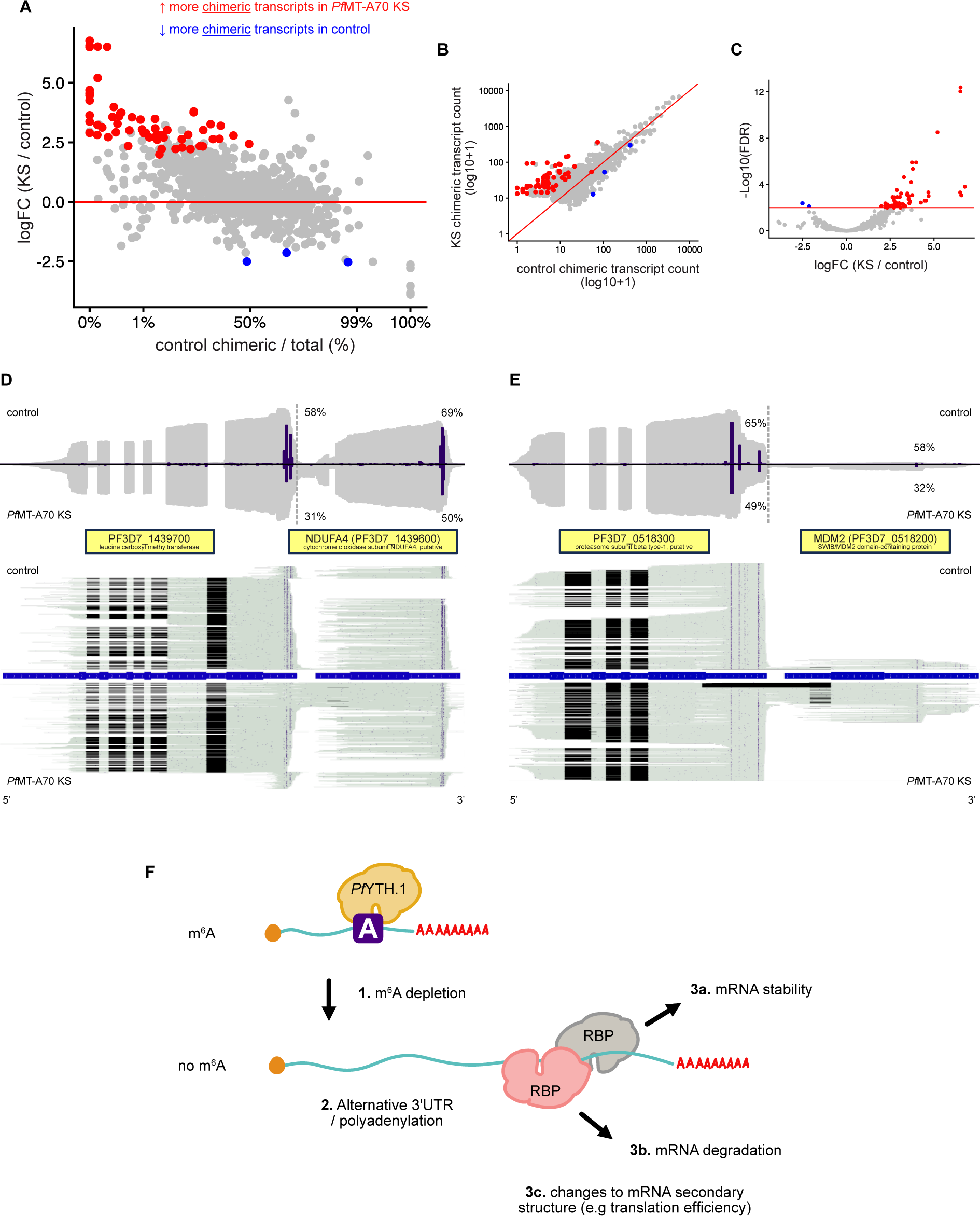
*Pf*MT-A70 mislocalisation causes more chimeric transcripts. **A)** Differential abundance of chimeric transcripts (those which map to multiple genes) shows an increase after *Pf*MT-A70-sandwich mislocalisation (28hpi, FDR < 0.01, GLM likelihood ratio test, edgeR interaction group vs chimeric). The interaction term highlights genes as significant when amount of chimeric transcripts change independently of background expression (total transcripts). **B)** Distribution of chimeric transcript count before and after *Pf*MT-A70 mislocalisation highlighting significantly DE’d genes from a). **C)** Volcano plot shows genes with a significant decrease in chimeric transcripts were close to probability threshold. **D)** Genes with significant changes in chimeric transcripts across all timepoints were found by identifying top contributors in PCA analysis. IGV screenshots and coverage plots show two top PC2 loading contributors. PF3D7_1439700 shows lower m^6^A levels in transcripts which also map to the 3’ downstream gene NDUFA4. **E)** PF3D7_0518300 shows alternative splicing in some of the chimeric transcripts after *Pf*MT-A70 knock-sideways, indicated by introns (black regions within transcripts). **F)** Hypothesis of how m^6^A depletion could alter mRNA abundance by revealing context specific regulatory elements in alternative 3’UTR’s.

We then attempted to find genes which had the most significant proportional increase in chimeric transcripts. We first conducted a PCA on the chimeric transcript counts and found samples separated by both timepoint and treatment (Supplementary Figure 8A). We looked at some of the most influential genes from this PCA in IGV and found they had complex methylation patterns near polyadenylation sites and significant increases in chimeric transcripts, of which some were alternatively spliced (Figure 7D, 7E). In summary, these analyses revealed that a loss of m6A in 3’UTRs caused a failure of normal transcript termination and polyadenylation and resulted in abnormally lengthened transcripts – sometimes this increased length was sufficient that transcripts encompassed an entire additional downstream gene.

## Discussion

Errors in the processes of transcription, splicing and polyadenylation can result in defective gene expression and hence may disrupt cell proliferation. At the same time, error-prone transcript processing can generate indels and alternative isoforms that may create multiple gene products that potentially allow selection for cells to rapidly adapt to changing environments. Here, we show that the m^6^A modification is enriched at sites ∼50 nt upstream from transcript termination and polyadenylation. Induced global depletion of that modification in mRNA results in the cellular machinery missing the normal termination site in many transcripts, resulting in transcriptional readthrough of nuclear mRNAs and transcriptome-wide differential expression in *P. falciparum*. Our study corroborates similar findings in *Toxoplasma* and *Arabidopsis*, and other studies on m^6^A-dependent UTR lengths, while exposing the complexity of 3’UTR methylation and its role in alternative polyadenylation (29, 30).

In this study we define the m^6^A methylation patterns at a base resolution in *P. falciparum* genes. Preferential 3’UTR distribution of m^6^A has been shown in other eukaryotes, but we show that m^6^A often occurs at a very biased position around ∼50 nt upstream of polyadenylation sites, even for genes with alternative polyadenylation. We found that within a single gene individual m^6^A sites are depleted unequally after methyltransferase disruption. Questions remain around what directs canonical methylation. In *Plasmodium* and in other organisms, m^6^A is preferentially deposited at DRACH motifs, yet the majority of DRACH motifs genome-wide and within 3’UTRs were not methylated. DRACH motifs themselves are clearly not sufficient to define a site that will be methylated. Conversely, we saw a high degree of methylation at non-DRACH motifs. Many of these were detected by both the ONT RNA-seq and GLORI-seq, reassuring us these are not the false positives of one individual method. Further classification of 3’UTR methylation patterns based on number of canonical m^6^As and degree of methylation is needed to uncover what directs methylation and therefore determines 3’UTR formation in *P. falciparum*. Distance to neighbouring genes, extended DRACH motifs, upstream/downstream contexts (RBP sites) and gene essentiality should also be considered. We speculate that in some genes, multiple m^6^A sites act as fallback termination site for polyadenylation machinery that has inefficient substrate recognition. We propose two potential explanations for this apparent sloppiness. One is that this inefficiency generates variation which is an advantage for a parasite which must rapidly adapt to a harsh environment and active immunological adversary. Hedging bets on alternative polyadenylation and UTRs may increase survival for some individuals in perturbed environments. Alternatively – previous evidence has suggested a relatively high error rate in various aspects of *Plasmodium* gene regulation – e.g. high degrees of apparently aberrant intron retention (39) and sloppiness in eliminating transcripts through nonsense mediated decay (40). It may be that *Plasmodium’*s intracellular parasitic lifestyle provides an excess of energy and raw ingredients for macromolecular synthesis that permits a high degree of wasteful production of non-functional transcripts. Future studies should compare the complexity of 3’UTR methylation patterns in *P. falciparum* against other organisms to understand if and how methylation complexity is related to taxa.

The transcriptional readthrough after *Pf*MT-A70-sandwich mislocalisation is likely a two-step effect. First, inducible mislocalisation of the catalytic methyltransferase subunit *Pf*MT-A70 from the parasite nucleus to plasma membrane results in decreased methyltransferase activity and therefore less m^6^A in mRNA. Second, depleted levels of m^6^A impair transcript recognition by CPSF – the enzyme complex responsible for endonucleolytic cleavage and recruitment of polyadenylation machinery to mRNA. In mammals, CPSF consists of at least 6 subunits (41), one of which shares homology to the *Plasmodium* m^6^A reader *Pf*YTH.1 (CPSF4 / CPSF30) (9). It follows that without normal levels of m^6^A in mRNA, *Pf*YTH.1 becomes defective in its direction of CPSF to canonical polyadenylation sites. The two CPSF subunits CPSF30 and Wdr33 bind directly to the polyadenylation signal (AAUAAA), which is well characterised in mammals, but poorly in *Plasmodium* (37, 42, 43). Given the AT-richness of the polyadenylation signal and *P. falciparum* UTRs (90% AT), it is possible that a mammalian-like polyadenylation signal is not sufficiently specific in *Plasmodium*, placing greater dependence on the m^6^A-*Pf*YTH.1 recruitment of CPSF for proper UTR processing. Another CPSF subunit, Fip1, guides polyadenylation by binding to U-rich regions in mRNA, an additional polyadenylation signal which is potentially muddied by the AT-richness of *Plasmodium* (44, 45). Additionally, it remains unknown which m^6^A sites detected in our RNA-seq were the result of co- or post-transcriptional methylation (46). To terminate transcription, at least one m^6^A would have to be deposited co-transcriptionally. While co-transcriptional m^6^A deposition appears important to prevent transcriptional readthrough we cannot exclude the possibility that m^6^A deposition and further cleavage events occur post-transcriptionally.

We found that this m^6^A dependent polyadenylation signal was present in most mRNAs, irrespective of gene function. Genes which were differentially expressed, differentially methylated, or had differential abundance of chimeric transcripts had no clear gene ontology enrichment. While the absence of the m^6^A methyltransferase causes readthroughs to most mRNAs, the effect is contextual. First, we consider the case where a ‘lonely’ gene (which has no close gene neighbours) loses its m^6^A and consequently gains a longer 3’UTR. It is well reported that 3’UTRs regulate gene expression, and we speculate that a m^6^A placement regulates gene expression by preventing transcriptional readthroughs and creating an aberrantly long transcript that might contain unwanted binding sites for *trans* factors (addition of RBP-domains) or *cis* (changes to secondary structure). mRNA fate would then depend on what regulatory elements are revealed by transcriptional readthrough, with the possibility of increased or decreased expression based on the surrounding genomic context. Second, for convergent gene neighbours, transcriptional readthrough could cause clashes in RNA polymerase II on opposing strands, reducing expression of both genes (47). Third, chimeric transcripts may or may not get translated into functional proteins with unpredictable and likely deleterious effects. On the other hand, the removal of m^6^A could result in a chimeric transcript which gets translated (though the versions we found generally contained interrupting stop codons), explaining one method by which m^6^A-*Pf*YTH.1 acts as a programmed gene repressor (as is potentially the case for *Pf*ALAD and *Pf*SPP (48)). In all cases, it is difficult to tease apart the effect of m^6^A on individual transcription regulation when depleting m^6^A levels globally. This could be further explored by gene- and site-specific m^6^A-editing to reduce confounders.

Our analysis of transcriptome-wide methylation showed the importance of considering canonical and low frequency m^6^A methylation. m^6^A abundance in mRNA in mammalian cells has been reported as approximately 0.5% m^6^A / A (49). A problem in estimating an overall percentage for m^6^A abundance is the difficulty in assigning true confidence values to prediction of low abundance m^6^A sites. A single m^6^A predicted in a position not shared in other reads might reflect a sporadic off-target methylation by the writer complex, or incorporation of an m^6^A base by the RNA polymerase during transcription, or a false positive detection through sequencing. Differential methylation analysis suggests the latter is unlikely to explain all these low-frequency sites, as they were less abundant after *Pf*MT-A70 mislocalisation. We observed this when considering overall m^6^A levels per gene (Figure 3A). Furthermore, *P. falciparum* has no purine synthesis and relies on import and salvage pathways (22, 50). Since *Plasmodium* depends on purine salvage more than other model eukaryotes, there is potential opportunity to study non-specific polymerase activity and how modified nucleosides are incorporated into mRNA, to better understand the mechanism by which low-frequency methylation occurs and its impact on gene expression.

Finally, we highlight the importance of using genome browsers (e.g. IGV) in inspecting novel splicing and methylation patterns in mRNA flagged by tools designed to flag differential expression or usage. By inspecting differentially expressed genes in IGV, we saw that transcriptional readthrough of UBC E2 was likely responsible for increased transcript mapping to a 3’ neighbour gene, something which would be difficult to find without alignment viewers (Supplementary Figure 8E). The same browser allowed us to see novel splicing across chimeric transcripts when confirming differential abundance of chimeric transcripts (Figure 7D). Just as microscopists show cellular localisation or morphology and quantify those results with fluorescence assays, we encourage inspecting RNA-seq in genome browsers to validate expression or RNA modifications generated from headless software.

This study provides base level resolution of m^6^A methylation in asexual blood stages of *P. falciparum,* demonstrating the complexity of m^6^A methylation and its role in transcript processing. We emphasise how visualisation of ONT RNA-seq proved essential to our analysis by allowing us to discover transcript specific modification patterns which may have been missed using traditional differential expression or methylation software. We speculate that m^6^A has more importance as a polyadenylation signal in an AT-rich genome where a mammalian-like polyadenylation signal becomes non-specific. This m^6^A-polyadenylation signal is near ubiquitous in *P. falciparum* nuclear mRNAs, and its removal through methyltransferase perturbation induces widespread gene expression changes and transcriptional readthrough.

## Data Availability

Two RNA-seq datasets were generated independently for this study. The ONT RNA-seq dataset was generated at Bio21, University of Melbourne by the Ralph lab and the approach to culturing, RNA isolation, sequencing, processing and analysis are described in full below. The GLORI-seq dataset was generated at Institute Pasteur by the Baumgarten lab and its methods are described in supplementary materials and methods.

All code used to generate figures (as described in the following sections) is publicly available at https://github.com/levend1s/01_m6A_3p_readthrough_analysis. Scripts for generating QC figures are uploaded under https://github.com/stevelan/pfal_m6a_qc/.

ONT RNA-seq is uploaded to Sequence Read Archive under BioProject ID PRJNA1391863. GLORI-seq is uploaded under PRJNA1443230.

## Materials and Methods

### Generation of *Pf*MT-A70-2xFKBP-GFP-2xFKBP strains

To generate a mislocalisable GFP tagged version of *Pf*MT-A70, the pSLI plasmid (32) was utilised. The pSLI construct contains the 3’ of the gene of interest positioned inframe and upstream of a GFP marker. The GFP itself is flanked by 2 FKBP domains on either side (for a total of 4 FKBP domains), and followed by a 2A skip peptide and a neomycin resistance marker. The 3’ 629 nucleotides of *Pf*MT-A70, lacking the stop codon, was amplified using the primers Forward: GCGGCCGCGGTCCTCAATGGATACGATGTG and Reverse: CCTAGGCGCGTTTGGATTATTTCTTGGG, with *NotI* and *AvrII* restriction sites respectively, and cloned into *NotI* and *AvrII* sites of the pSLI plasmid. This was transfected, parasites and integrated transfectants selected as described by Birnbaum and colleagues (32). A pLyn-FRB-mCherry mislocaliser (containing a plasma membrane anchor) was subsequently transfected into this strain, and double transfectants selected as described previously (32). Parasites transfected with both plasmids were verified by fluorescence microscopy for expression of both GFP and mCherry.

### Parasite culture

*P. falciparum* parasites were cultured in human O+ erythrocytes kindly provided by the Australian Red Cross Lifeblood with an adapted method described previously (51). Infected erythrocytes were cultured at 5% haematocrit in RPMI 1640 (Gibco # 31800105) supplemented with 25 mM HEPES (Formedium # HEPES05), 0.5% (w/v) AlbuMAX™ II (Gibco # 11021045), 13 μM gentamycin (Enzo # ALX380003G005) and 140 μM hypoxanthine (Sigma-Aldrich # H937725G) at 7.4 pH (hereafter referred to as media). Parasite cultures were maintained in a sealed Perspex box filled with malaria gas mix (1% O_2_, 5% CO_2_, 94% N_2_) and stored at 37°C to mimic in vivo conditions. Parasitemia was monitored in cultures by routinely inspecting Giemsa-stained blood smears.

Parasites were synchronised to a 4-hour window by magnetic purification of schizonts followed by sorbitol treatment. Briefly, resuspended culture was flowed through a CS column (Miltenyi Biotec) held in a magnetic apparatus. The CS column was flushed with media to ensure only infected RBCs (iRBCs) containing late-stage parasites remained in the column. The column was then removed from the magnetic apparatus and flushed with media capturing the flow through, to obtain the magnetically captured iRBCs. Reinvasion occurred on an orbital shaker at 37°C for four hours followed by 5% (w/v) D-sorbitol treatment (Sigma # S3889) (52), marking the four hpi timepoint.

100 nM rapamycin (Cayman # 13346) was added to knock-sideways replicates four hours prior to each trophozoite harvest and culture was resuspended. Parasites were then harvested by treatment with 0.03% (w/v) saponin in duplicate as control or knock-sideways at 28, 32 and 36 hpi (12 samples total).

### Growth assay

*Pf*MT-A70 knock sideways parasites or *Pf*3D7 were synchronised to a 2-hour window by Percoll (Cytiva # 17089102) gradient purification of late-stage parasites, followed by sorbitol selection of early-stage parasites. High parasitaemia cultures were layered on a 65% Percoll in 1×PBS solution and centrifuged. The late-stage layer was transferred to a dish with fresh media and RBCs and allowed to invade with gentle agitation. Newly invaded parasites were selected using 5% (w/v) D-sorbitol. The parasitaemia was adjusted to 0.2%. 100 nM rapamycin was added to the *Pf*MT-A70 knock sideways group (n=3) and *Pf*3D7 rapamycin group (n=2) and controls (n=2 for each) were treated with DMSO. The parasitaemia was measured using Giemsa-stained thin smears at 30 hpi and 48 hours following this timepoint to measure the growth rate.

### Fluorescence microscopy

Samples were imaged using a DeltaVision Elite Widefield Deconvolution microscope (General Electric Healthcare). Slides were imaged with TRITC and FITC filter sets for emission and excitation wavelengths. Processing of deconvolved images was done with FIJI ImageJ software (v2.9.0).

### Total RNA isolation

Cultures were centrifuged (500 rcf for 5 minutes) and resuspended in 10X sample volume of 0.03% (w/v) saponin (Sigma # S4521) in 1X PBS before incubating at room temperature for 10 minutes and centrifugation (1800 rcf for 5 minutes at 4°C). Parasite pellet was washed in 1 mL 1X the sample volume of PBS and centrifuged (10,000 rcf for 30 seconds) before resuspending in 1 mL TRIzol™ (Invitrogen # 15596018).

RNA was extracted from samples suspended in TRIzol™ using a modified protocol based on the TRIzol™ reagent user guide. Briefly, 0.2X the sample volume of chloroform was added to the TRIzol™ sample, then vortexed and incubated at room temperature for 3 minutes. Samples were centrifuged (12000 rcf for 30 minutes at 4°C) to induce phase separation and the upper phase (RNA) for each sample was transferred to a clean Eppendorf tube with 1X sample volume of 80% absolute ethanol in RNase-free water. The 36 hpi samples did not correctly phase separate so were vortexed and centrifuged again, whereafter they successfully phase separated.

Precipitated RNA was purified according to the RNeasy® MinElute® Cleanup column protocol (QIAGEN # 74204). DNase I digestion of eluted RNA sample was carried out to remove residual DNA, before performing a second RNeasy® MinElute® column cleanup. During the second MinElute® cleanup, the final RNA sample was eluted from the column in two elutions of 60 μL RNase-free water. Ethanol precipitation was carried out to increase concentration as required by Nanopore SQK-RNA004 flow cells.

Integrity of RNA samples was verified using Agilent 2200 TapeStation electrophoresis system according to the manufactures protocol ‘Agilent RNA ScreenTape System Quick Guide’. Concentration of RNA samples were evaluated using an Invitrogen Qubit 3 according to the manufacturers protocol ‘Qubit RNA High Sensitivity Assay Kit’.

### RNA sequencing

An ONT hybrid RNA-cDNA library for sequencing was created according to the manufacturers protocol ‘Direct RNA sequencing SQK-RNA004’. (NB in this procedure the synthesis of the complementary DNA strand provides stability to the RNA strand, but it is the native RNA strand that is sequenced). Sequencing was performed on ONT MinION Mk1B using FLO-MIN004RA flow cells connected iMac with an M2 chip ram running MinKNOW (v24.06.5) and basecalled using the rna004_130bps_fast@v3.0.1 model. Samples were re-basecalled on the Melbourne University Spartan HPC platform with Dorado (v0.8.3) using the rna004_130bps_sup@v5.1.0 model.

### RNA-seq alignment and filtering

Generated modBAMs were aligned to reference genomes using Dorado, which uses Minimap2 and uses the Minimap2 ‘lr:hq’ preset, which is optimised for aligning long reads produced by Nanopore RNA-seq (53). Reference genomes for *Saccharomyces cerevisiae* and *Homo sapiens* GRCh38 were obtained from Ensembl (54). The reference genome for *P. falciparum* 3D7 (release 68) was obtained from PlasmoDB (55). Separate alignment files were generated for each reference genome. Reads which mapped to *H. sapiens* with MAPQ ≥ 1 and reads which did not map to the *Plasmodium* genome were removed from RNA-seq files prior to SRA upload. RNA-seq analysis of *P. falciparum* used a filtered BAM file generated with SAMtools filtered for high mapping quality (MAPQ ≥ 50) and removing any reads which mapped to the human genome (56). Gene counts for aligned reads were generated using the featureCounts program (v8.3) from the Subread package with long read, overlap and strandedness options (38).

### RNA-seq quality control

Read length statistics were generated with Cramino (v1.2.0) producing columnar, arrow file format statistics that were then summarised with NanoPlot (v1.46.1).

Unaligned, unfiltered and MAPQ > 50 filtered bam files were inspected using pysam (v0.23.3). Reads were stratified by mean quality score into strata of size five. Mean quality scores for reads were calculated by first converting each bases Q score to an error rate, averaging the error rates and then converting the average back to PHRED Q scores. Nanopore produces probabilities scores as values between 0 and 255. The distribution of modification probabilities were binned with every bin containing 4 modification probabilities and then plotted as a histogram, values Q25 and above were all treated as the one bin. Only m^6^A modification probabilities > 0.5 were considered.

Feature counts were calculated for all RNA features in the *P. falciparum* reference gene feature format (gff) file. A feature was counted as long as a single base from a read overlapped its gff co-ordinates. This categorised reads which overlapped RNA features into mRNA, rRNA, ncRNA. Plots were generated comparing unfiltered reads to filtered reads. Reads filtered by alignment score increased the proportion of mRNA. Samples were pooled by timepoint for hatched bar graph and pooled across all samples for RNA feature type pie chart.

### Differential transcript abundance analysis

Differential expression analysis was performed using the likelihood ratio test GLM in edgeR (v4.4.2) (57) and visualised with ggplot2 (v3.5.2) (58). Lowly expressed genes (< 10) were filtered. An FDR threshold of 0.05 was used to define significance in expression change.

### Modification quantification

Bedmethyl files were generated using ONT Modkit (v0.4.2) using a modification probability threshold of 0.95 to generate genomic pileup information and study modifications in modBAMs files. Transcript coverage was generated using rqc.py. Transcriptome-wide modification abundance was generated using modkit summary. Reads were visualised with Integrative Genome Viewer (IGV) (v2.19.6) (59).

Site-specific differential methylation based on treatment was performed using the negative binomial GLM and methylation status as an interaction term in edgeR. Sites were first separated into nuclear or mitochondrial groups and processed separately. Filtered sites had ≥ 20 m^6^A, were not differentially expressed (negative binomial GLM) and were not in genes whose annotation overlapped by at least 1 nt. Gene type and location (UTR vs CDS) was mapped to each site using bedtools (v2.31.1) (60).

Weighted average methylation for canonical (m^6^A / A ≥ 50%), low m^6^A frequency (m^6^A / A < 50%) and total methylation were calculated for each gene using the scripts available at https://github.com/levend1s/rqc and visualised with ggplot2.

Canonical m^6^As were located and mapped to genes with rqc.py gene_methylation_analysis. This output, along with the estimated canonical polyadenylation from approximate_tes, were used to generate canonical m^6^A offset plots using rqc.py ‘calculate_offsets’ and ‘plot_relative_offset’ functions. Histograms of transcript end sites separated by canonical m^6^A were generated using rqc.py m6A_specific_tes_analysis function, filtering for transcripts with a poly-A tail length ≥ 10 nt, genes with read depth ≥ 10, and by manually specifying a plot offset as the approximated canonical polyadenylation site.

In our analysis we observed ‘m^6^A smearing’, whereby homopolymeric runs of adenosine at the 5’ end of a DRACH-like motif (AC) saw many m^6^A detected within the homopolymer. This is likely from a technical error from ONT difficulties in homopolymeric basecalling, and we observed deletions were exacerbated in similar contexts. These smears, if false positives, would affect average differential methylation results. Poor annotations also invite inaccuracies in calculating canonical methylation as sites may be missed or incorrectly associated with a gene. Calculating average of differential methylation is somewhat compromised by complex methylation patterns, nanopore errors, and poor annotations and these should be considered when performing differential methylation analysis.

### Chimeric transcript and readthrough analysis

Chimeric transcripts were identified using featureCounts with the overlap option and the default overlap size of 1 nt. This produced a file with each RNA-seq read ID and how many genes (features) it mapped against. This file was processed (with bash text processing - awk, uniq, sort, tr) to find for each gene the count of reads which mapped that gene and at least one other (chimeric).

Differential abundance of chimeric transcripts was performed using the negative binomial GLM in edgeR and visualised with ggplot2. Genes with total 0 chimeric transcripts across all samples at a timepoint were filtered out. An FDR threshold of 0.05 was used to define significance in expression change. Model used treatments (control vs knock-sideways) as groups with chimeric transcript vs non-chimeric transcripts as an interaction term, to discover genes with significant proportional changes of chimeric transcripts.

PCA was done on chimeric transcript count matrix using prcomp from the R stats package. Scatter plot of the loadings of each gene for PC1 and PC2 were plotted, with size and colour as a linear combination of the loading for each component. Top 20 loadings (linear combination) were labelled. Heatmap for the top 50 loadings for each component were plotted with colour as a normalised z-score of transcript expression with pheatmap (v1.0.13). Alignments of chimeric transcripts were imaged with IGV and coverage plots generated with rqc.py.

Canonical methylation and readthrough correlation plot used values from rqc gene_methylation_analysis and approximate_tes. Gene neighbour map was created using rqc.py gene_neighbour_analysis and used to filter gene sets in the correlation plots.

Readthrough proportional change was found by estimating a canonical polyadenylation site for each gene. This filtered for transcripts which had a poly-A tail ≥ 10 nt, and genes with read depth ≥ 10. Canonical polyadenylation was the highest peak (depth ≥ 20) in the histogram for each genes collection of transcription end sites, determined using scipy.signal.findpeaks with a minimum distance between peaks of 50 nt. An additional metric was reported called the apa_score, which was the ratio of the top two most prominent canonical polyadenylation site, and informs on the degree of alternative polyadenylation in a gene. After identifying canonical polyadenylation sites, readthrough proportion was calculated as the proportion of transcripts which terminated 3’ of the canonical polyadenylation site before and after treatment. In this way, readthrough proportion, average 3’UTR and poly-A tail length were calculated with the rqc.py approximate_tes function. These values averaged across samples against library size and plotted using ggplot2.

### ONT dRNA-seq and GLORI-seq overlap analysis

Sites common to ONT and GLORI were identified with bash text processing (‘awk’, ‘sort’, ‘uniq’) and bedtools ‘intersect’ (60). Bedmethyl and GLORI-seq files were pre-processed to keep only sites with m^6^A / A ≥ 25% and read depth ≥ 15. DRACH sites were identified using rqc.py motif_finder command. Generated BED files were imaged using IGV (v2.19.6) against ONT RNA-seq (59). Gene body-coverage plots were generated with these bed files using rqc.py plot_coverage with options subfeature_cds, 100 bins and 500 nt coverage padding.

### GLORI-seq - Parasite culture

Asexual blood-stage wildtype 3D7 *Plasmodium falciparum* parasites were cultured in human red blood cells (obtained from the Etablissement Francais du Sang with approval number HS 2021-24819) in RPMI-1640 medium (Thermo Fisher # 53400-025). The medium was supplemented with 0.5% w/v AlbuMAX™ I (Thermo Fisher no. 11020039), hypoxanthine (0.1 mM final concentration, CC-Pro # Z-41-M) and 10 mg gentamicin (Sigma # G1397- 10 mL) at 4% haematocrit and under 5% O2, 3% CO_2_ at 37°C. The parasite development was monitored by blood smears and Giemsa staining.

### GLORI-seq - Synchronization

To collect highly synchronous asexual stages, parasites were synchronized to a 6-hour window by Plasmion (Fresenius Kabi) treatment followed by sorbitol treatment. The 0-hour timepoint was considered to be 3 hours after the Plasmion treatment. The parasites were sampled in technical duplicates (total RNA isolation) 12 hours, 24 hours and 48 hours post invasion of the red blood cells.

### GLORI-seq - Total RNA and mRNA isolation

Red blood cells were lysed by 30 sec incubation with 0.075% Saponin in DPBS at 37°C followed by centrifugation at 4000 rpm for 5 min. The pellet was washed with ice-cold DPBS and resuspended in 700 μL QIAzol Lysis Reagent (Qiagen #79306). The total RNA was extracted with the miRNeasy® Mini kit (Qiagen #217004) according to the manufacturer’s protocol and subsequently selected for mRNAs with the Dynabeads mRNA DIRECT Kit (Thermo Fisher # 61011).

### GLORI-seq - GLORI sequencing and analysis

The GLORI sequencing protocol was adapted from Liu et al., 2023 (36) for *P. falciparum* as previously described (31). In brief, total RNA and mRNA were extracted from wildtype *P. falciparum* parasites at 12 hpi, 24 hpi and 36 hpi as described above. mRNA was fragmented at 94°C for 3 min in NEB RNA 10X fragmentation buffer (NEB # E6186A) and cleaned up by ethanol precipitation and the RNA Clean & Concentrator -5 kit (Zymo Research # R1014). Subsequently, the RNA (10 μL) was mixed with 6 μL glyoxal, 20 μL DMSO and 4 μL RNase-free water and incubated for 30 min at 50°C in a thermocycler (lid set to 100°C) for RNA protection. Following the incubation, 10 μL saturated H_3_BO_3_ were added, incubated for 30 min at 50°C and placed on ice for 2 min. For the deamination of the RNA, 50 μL of deamination buffer (5 M NaNO_2_, 500 mM MES (pH 6.0), 8.8M glyoxal) were added to the sample and incubated in a thermocycler for 8 hours at 16 °C and then hold at 4°C overnight. Afterwards, the RNA was precipitated as described above and the pellet was resuspended in 50 <u>μL</u> deprotection buffer (10 mL 1 M Triethylammonium pH 8.6, 9.5 mL deionized formamide, 0.5 mL RNase-free water) and incubated in a thermocycler at 95°C for 10 min. The RNA was again precipitated and cleaned as described above. For end repair, the reaction RNA from the previous step was mixed with 10X T4 PNK Buffer (NEB # B0201S), T4 PNK (NEB # M0201S), Murine RNase inhibitor (NEB # 0314S) and RNase-free water and incubated at 37°C for 30 min (lid set to 50°C) for 30 min. After addition of 10 mM ATP, the reaction was incubated for another 30 min. Next, RNA was cleaned once again with the RNA Clean & Concentrator -5 kit (Zymo Research # R1014). SmallRNA libraries were generated with NEBNext (NEB # E7300S). The small RNA libraries were pooled and size-selected using a Pippin Prep (Sage Science #CDF2010) and sequenced on an Illumina NextSeq 2000 in a 2×150 bp layout.

Using bcl2fastq2 (version 2.20, Illumina), raw reads were demultiplexed. The sequencing adapters were removed using trimmomatic (61). Using the GLORI-tools, STAR reference indices for a genome assembly featuring A-G conversion and a reverse complement of the latter were generated (36). Subsequently, STAR (62) was used to align the pre-processed sequencing reads to the A-G converted reference genome, allowing only uniquely mapping reads (option --outFilterMultimapNmax 1) Using the STAR default options, allowing only for uniquely mapping reads (option --outFilterMultimapNmax 1), reads that did not map to the A-G converted reference were aligned to the reverse-complement reference genome, too. The md flag was added to the BAM files by samtools’ ‘calmd’ after removing PCR duplicates with samtools’ ‘rmdup’ (56). JACUSA2 ‘call-1’ (63) was used to calculate the coverage of each nucleotide for each genome position. G references sites featuring a minimum coverage of 15 and a A-G conversion of >= 25% were retained as putative methylation sites since m6A sites are resistant to chemical A-G conversion. The overlap between replicates at each timepoint was determined using bedtools ‘intersect’ (60).

## Acknowledgements

This work was funded in part through a Discovery Project grant to SAR from the Australian Research Council (Grant number: 108083). We thank Traude H. Beilharz (Monash University) and Mike Clark (The University of Melbourne) for very helpful discussions on m6A and polyadenylation. We are grateful to Tobias Spielmann (Bernhard Nocht Institute) for the provision of inducible mislocaliser plasmids.

## Supplementary Figure legends

**Supplementary Figure 1. Nanopore assigns probabilities for base modification**

**A)** Plots show probability distribution for individually sequenced m^6^A sites. Distributions show a large spike in amount of m^6^A sites with high probability (≥ 0.95). This threshold is used in downstream analysis. **B)** IGV screenshots show modification sites in example gene under different thresholds (purple = m^6^A, red = m^5^C, green = inosine, yellow = pseudouridine). In this example, canonical m^6^A sites are largely unaffected by the threshold change while non-canonical m^6^A sites mostly disappear.

**Supplementary Figure 2. Analysis of driving genes in transcript differential expression analysis**

**A)** PCA based off gene mapped transcript count separates by timepoint, but not group. **B)** Gene set enrichment analysis (GSEA) for principal component 1 shows 23 GO enriched, including schizogony and cell growth terms. **C)** Normalised expression heatmap of top 50 loadings for PC1 show rhoptry and merozoite genes drive the principal component that describes timepoints, alongside numerous genes with unknown function.

**Supplementary Figure 3. Nuclear genome site level differential methylation analysis**

**A)** Site level differential methylation analysis shows sites with low initial methylation are more likely to increase in methylation after *Pf*MT-A70 knock-sideways. **B)** Sites within non-coding RNA are generally more lowly methylated than mRNA sites. **C)** From observation in IGV, unmapped sites tend to be in the 3’ region of transcripts from poorly annotated genes.

**Supplementary Figure 4. Weighted average methylation analysis**

**A)** PCA based off weighted average methylation (WAM) score separates by group, and weakly by timepoint. **B)** Gene set enrichment analysis (GSEA) for principal component 1 shows 9 GO terms enriched. **C)** Normalised WAM heatmap of top 50 loadings for PC1. **D)** Example WAM scores for genes with multiple canonical m^6^A. **E)** WAM change after *Pf*MT-A70 knock-sideways results is similar across genes with differing counts of canonical m^6^A. **F)** Genes with a greater read depth tend to have more canonical m^6^A. This could be explained by more transcripts leading to more alternative polyadenylation sites which each require a unique m^6^A site. **G)** Comparing WAM and read depth shows no correlation. This is desired and shows there is no read depth bias in WAM calculations.

**Supplementary Figure 5. Effect of MAPQ filtering on read count, RNA type and average base quality distribution**

**A)** Bar plot shows read count is reduced by approximately 50% after removing low quality reads (MAPQ ≥ 50), count for each group pools replicates (n=2) **B)** Pie charts shows majority of RNA types detected were mRNA, followed by rRNA and other RNA types (snRNA, snoRNA, tRNA, ncRNA). Proportion of rRNA reads is reduced by 12% after filtering reads for mapping quality. **C)** NanoPlot bivariate plot shows read length and average base quality (PHRED) with axis histograms, all samples pooled. Red arrows indicate the presence of *S. cerevisiae* enolase II control from RNA004 kit, and population of low PHRED reads which are absent after filtering for reads with high mapping quality (MAPQ) to *P. falciparum* genome.

**Supplementary Figure 6. Positional modifications in non-mRNA transcripts**

**A)** Cropped IGV screenshot of tRNA transcripts show detection of a Ψ (yellow) at the conserved TΨCG loop. **B)** Cropped IGV screenshot of *P. falciparum* 18S rRNA transcripts (PF3D7_0531600, A2) shows conserved positional modification of Ψ (yellow), m^5^C (red) and m^6^A (purple). Annotations show coordinates of known modifications in *Homo sapiens* (X03205) reference after blastn alignment of the two 18S genes. Note that some reference modifications differ from the detected modification: m^1^acp^3^Ψ (1-methyl-3-α-amino-α-carboxyl-propyl pseudouridine) is detected as Ψ, Cm (2′-O-methylcytidine) is detected as m^5^C, and m^6^_2_A is detected as m^6^A. **C)** IGV screenshot of entire mitochondrial genome (-ve strand genes only) shows positional modifications for Ψ and m^5^C.

**Supplementary Figure 7. Genome wide comparisons of WAM and run-on transcripts show weak correlation**

**A)** Average UTR length increases after *Pf*MT-A70 knock-sideways. **B)** Proportion of transcripts which terminate after the canonical polyadenylation site increases after *Pf*MT-A70 knock-sideways. **C)** Poly-A length decreases after knock-sideways for 28hpi and 32hpi, but increases at 36hpi. **D)** Comparing change in run-on proportion with WAM change shows a weak correlation. Often, genes have multiple m^6^A or close neighbours which may skew the run-on or WAM calculations. When excluding these genes by considering genes with a single m^6^A and no neighbouring genes, we still observe a weak correlation (28hpi, m^6^A / A ≥ 50%, depth ≥ 10). **E)** IGV screenshot of example neighbouring genes, where run-on transcripts from the 5’ neighbour could be counted towards the 3’ neighbour expression level.

**Supplementary Figure 8. Chimeric transcript analysis**

**A)** PCA based off chimeric transcript count (reads which map to multiple genes) separates by both timepoint and group. **B)** Scatterplot of genes based on their contribution to either principal component. **C)** Normalised WAM heatmap of top 50 loadings for PC1 and **D)** PC2. **E)** IGV screenshot of example gene PF3D7_1204000 which had 40-60x fold coverage increase after knock-sideways. There were at least two transcripts (highlighted red) sequenced which were spliced for both the 5’ upstream gene UBC E2 and PF3D7_1204000, and which lacked the canonical m^6^A present at the 3’ ends of UBC E2 transcripts, however these transcripts also mapped to the human genome and so were filtered from the uploaded RNA-seq.

