## Supplementary Figures for "m^6^A positions polyadenylation in *Plasmodium falciparum*"

### Supp figure 1

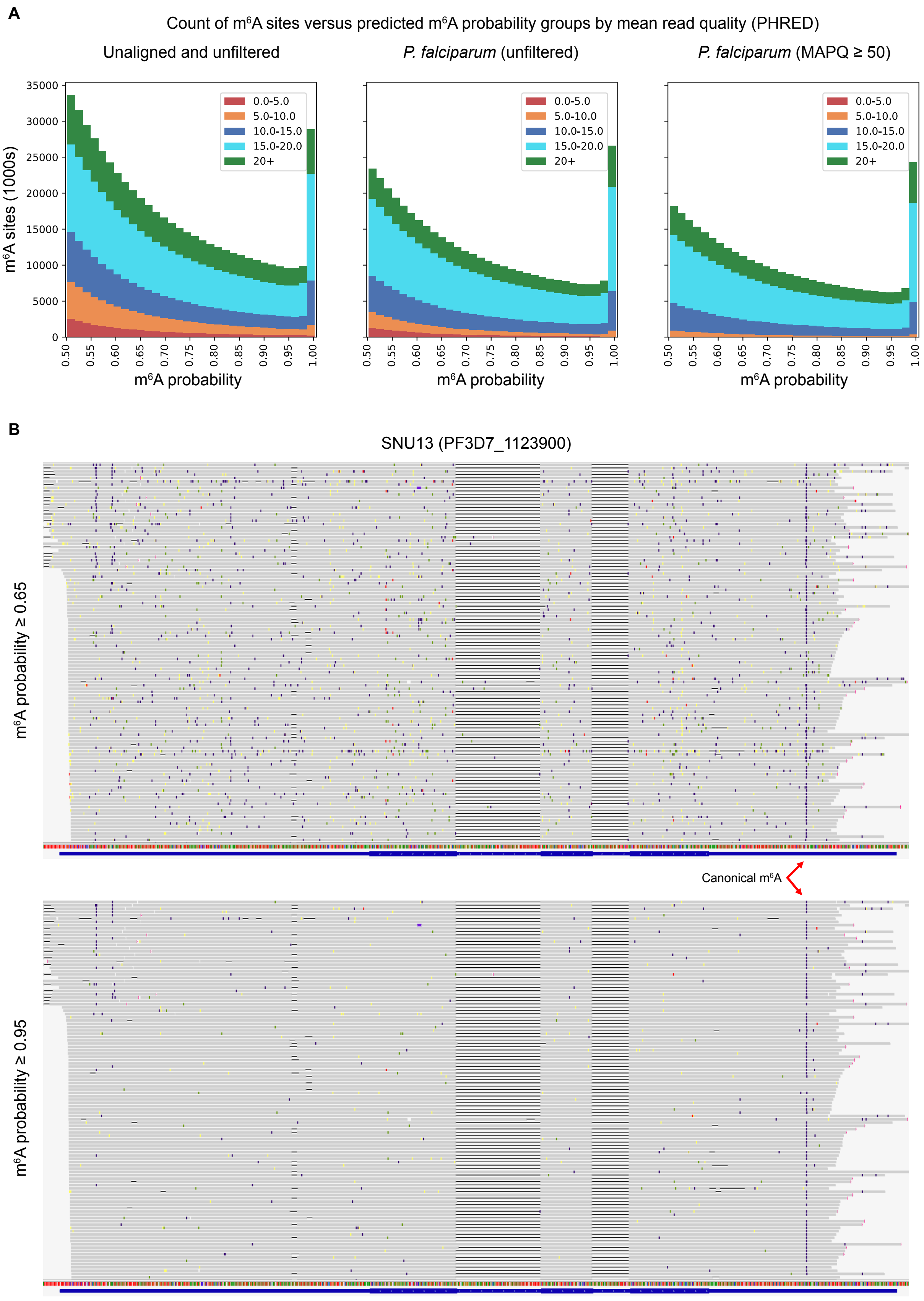

### Supp figure 2

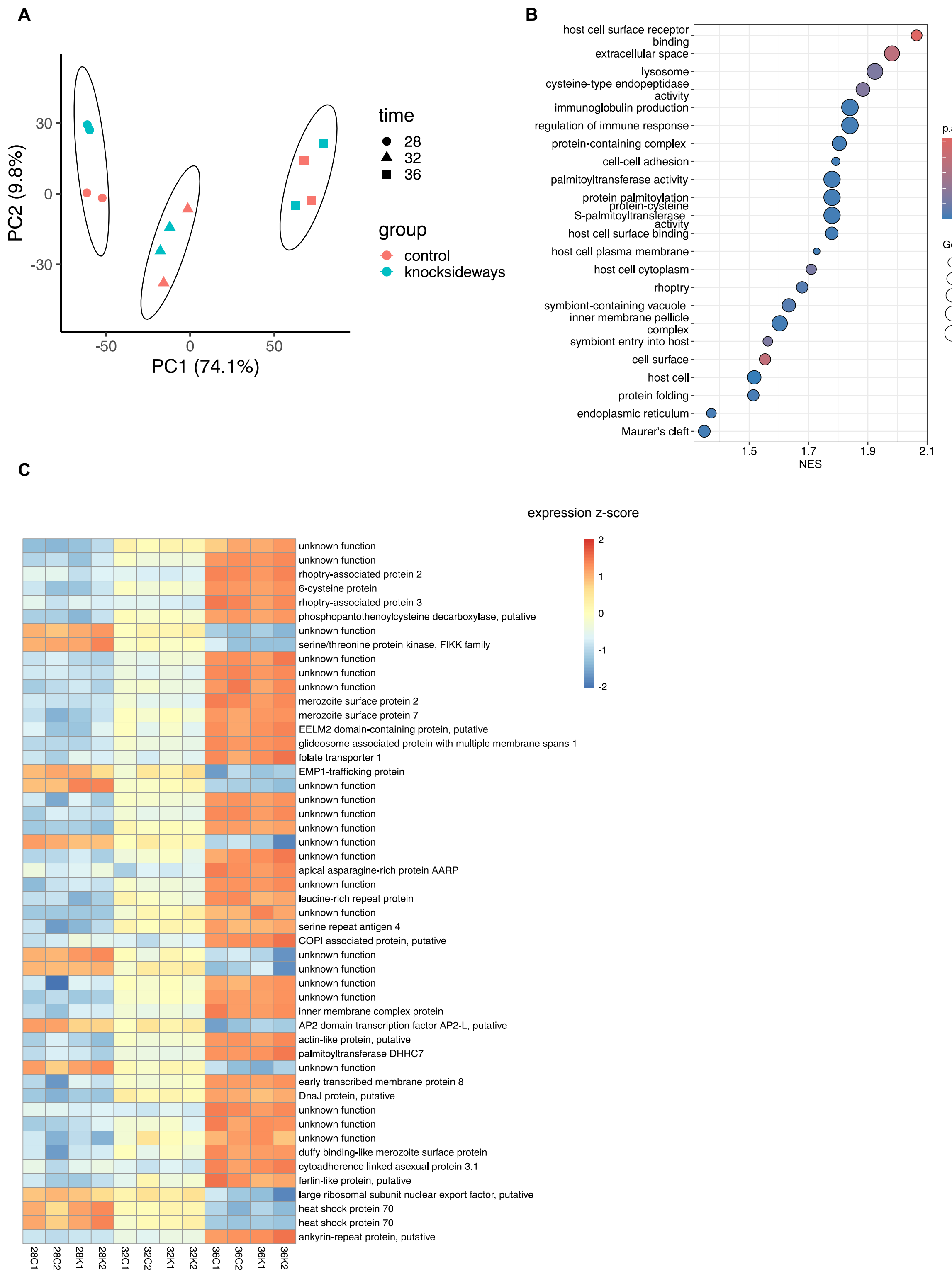

### Supp figure 3

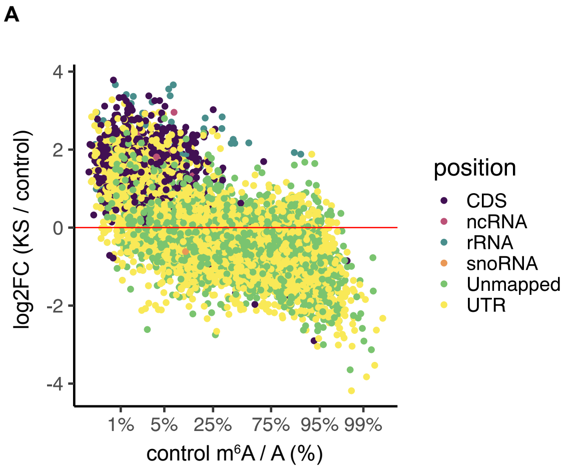


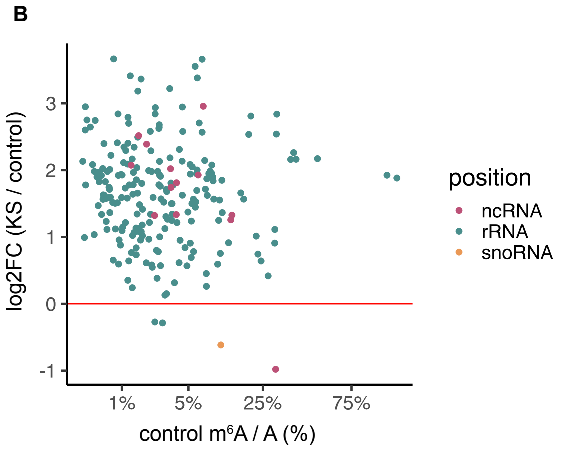


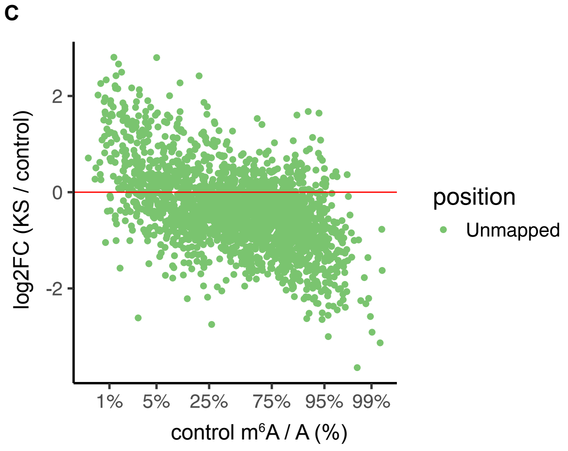

### Supp figure 4

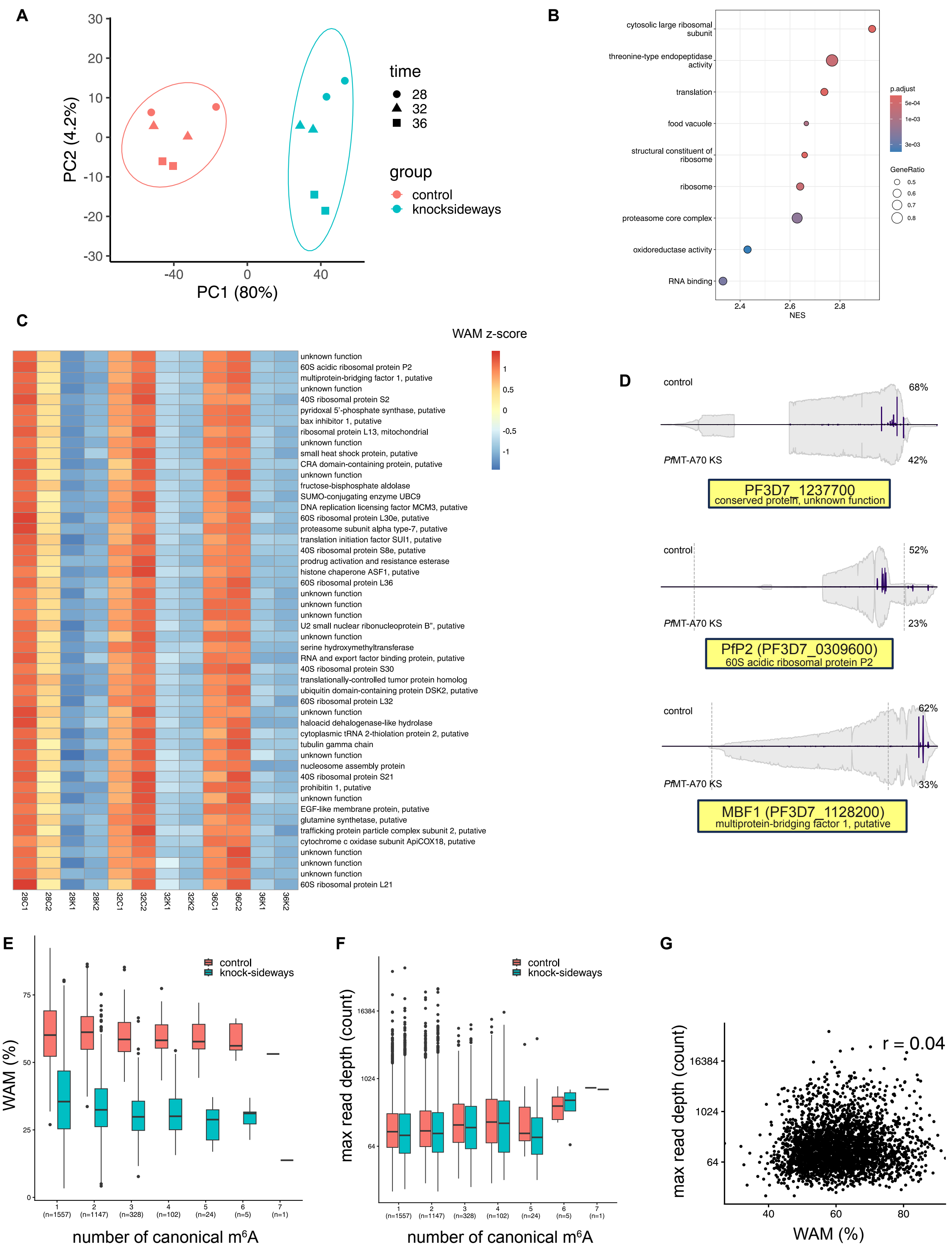

### Supp figure 5

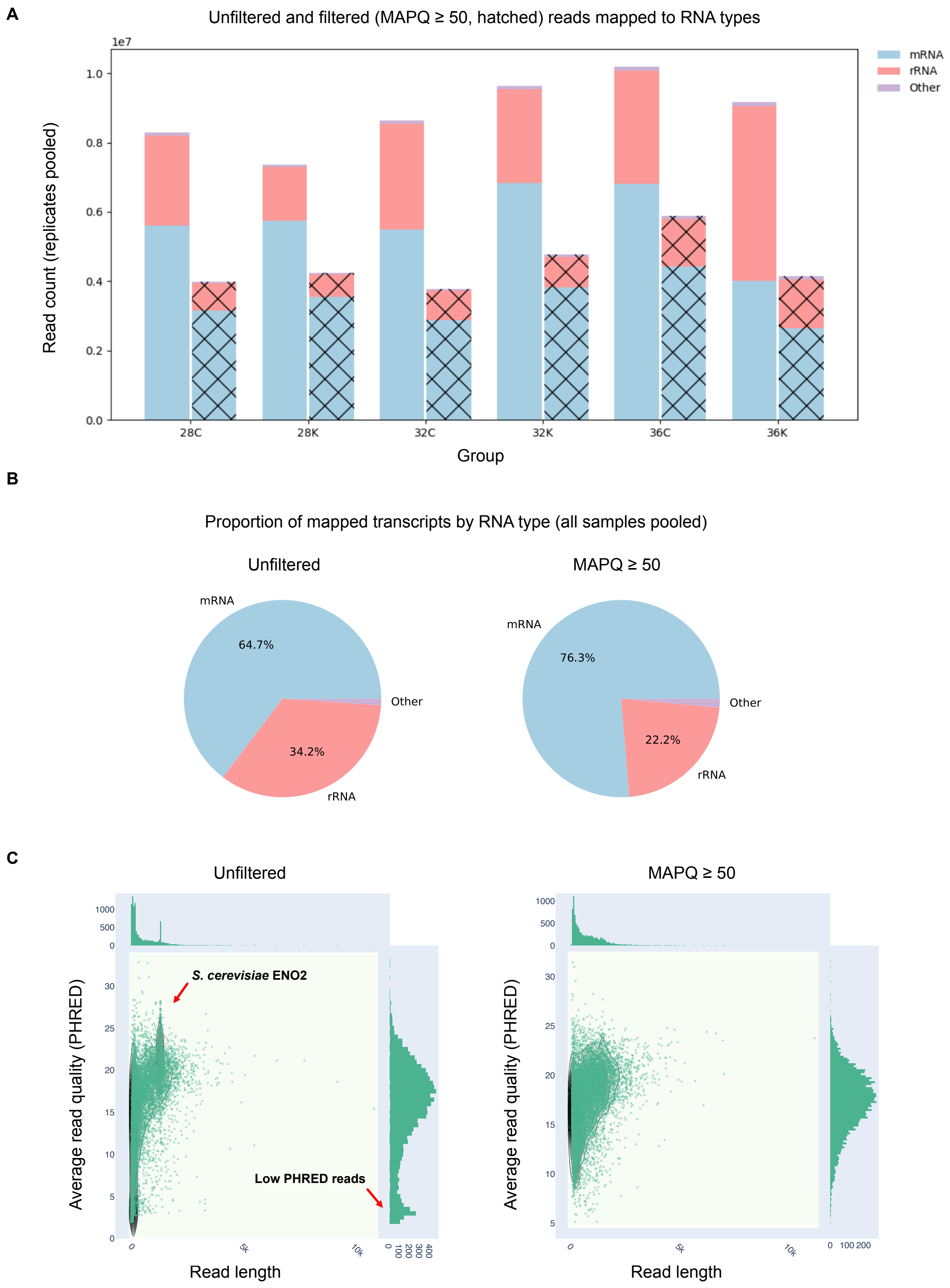

### Supp figure 6

A

**tRNA alignments**

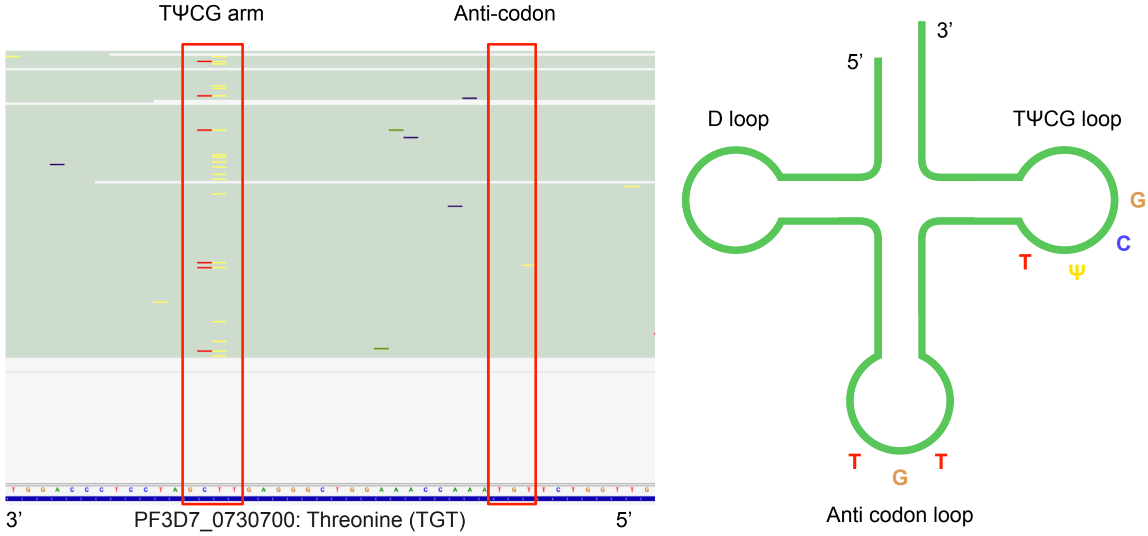


B

**rRNA alignments**

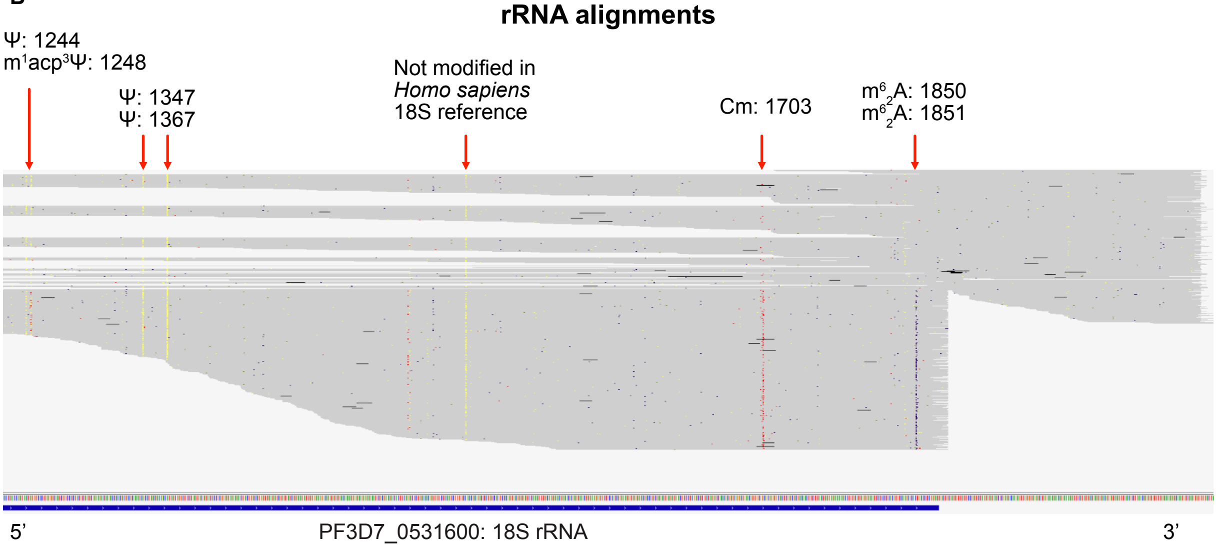


C

**Mitochondrial genome alignments (-ve strand)**

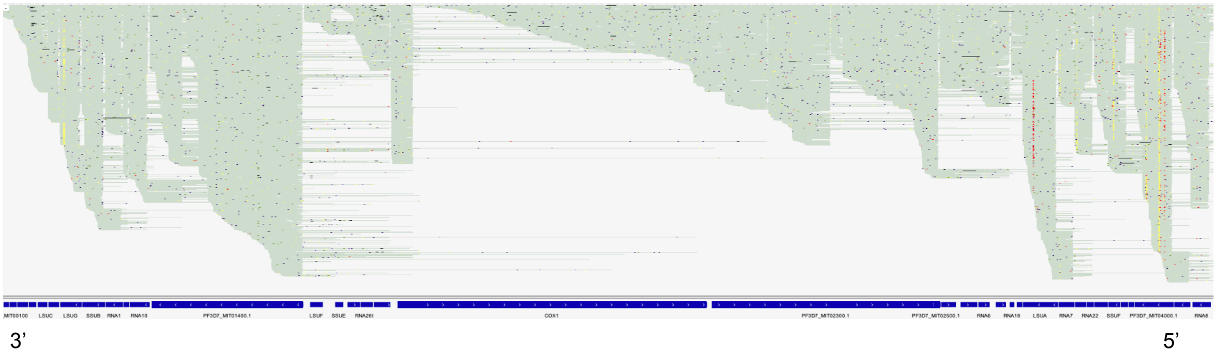

### Supp figure 7

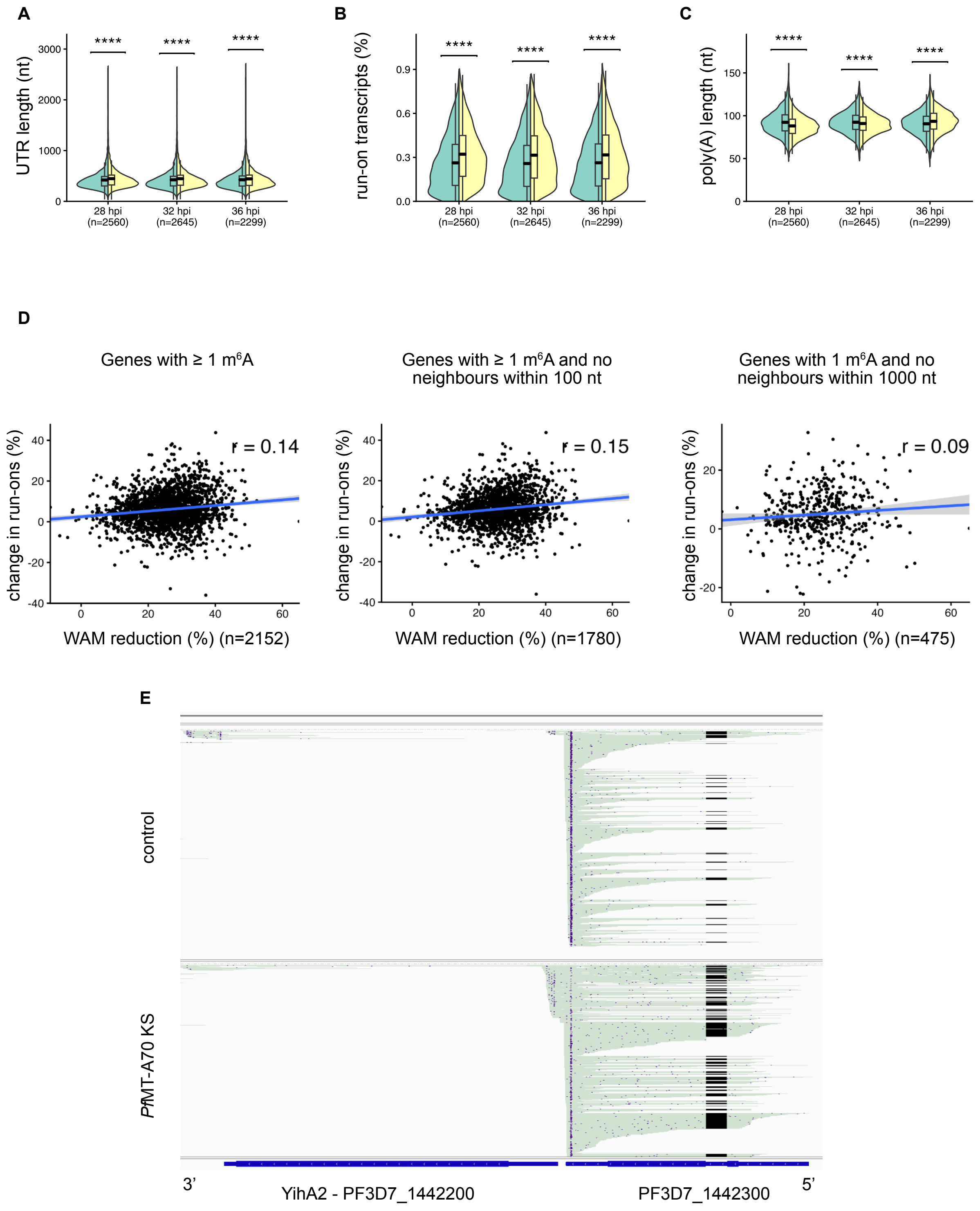

### Supp figure 8

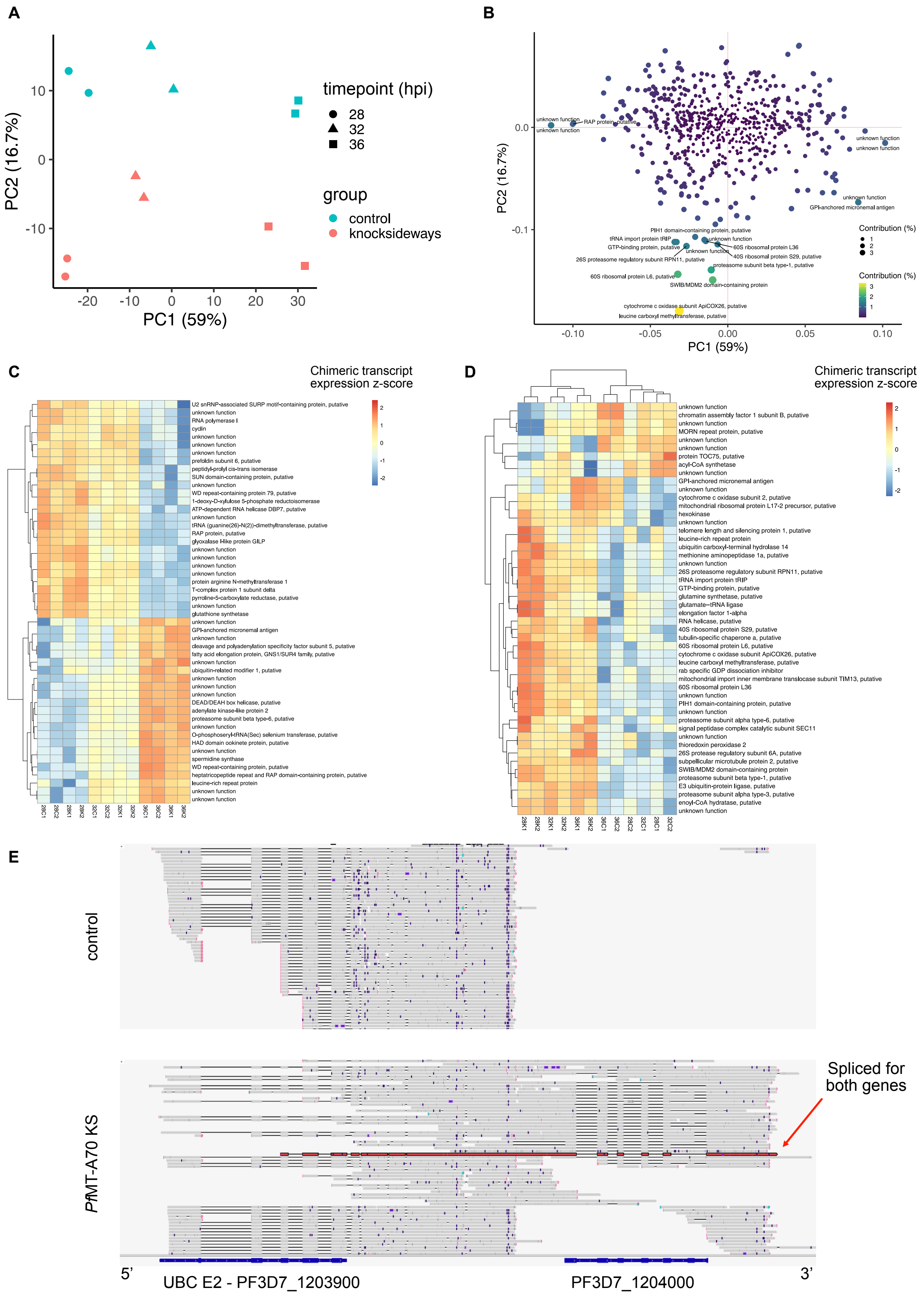
